# Quantifying the adaptive landscape of commensal gut bacteria using high-resolution lineage tracking

**DOI:** 10.1101/2022.05.13.491573

**Authors:** Daniel P.G.H. Wong, Benjamin H. Good

**Author notes:** Correspondence should be addressed to: B.H.G.

## Abstract

Gut microbiota can adapt to their host environment by rapidly acquiring new mutations. However, the dynamics of this process are difficult to characterize in dominant gut species in their complex *in vivo* environment. Here we show that the fine-scale dynamics of genome-wide transposon libraries can enable quantitative inferences of these *in vivo* evolutionary forces. By analyzing >400,000 lineages across four human *Bacteroides* strains in gnotobiotic mice, we observed positive selection on thousands of previously hidden mutations – most of which were unrelated to their original gene knockouts. The spectrum of fitness benefits varied between species, and displayed diverse tradeoffs over time and in different dietary conditions, enabling inferences of their underlying function. These results suggest that within-host adaptations arise from an intense competition between numerous contending mutations, which can strongly influence their emergent evolutionary tradeoffs.

## Introduction

The mammalian gut is home to a diverse microbial community comprising hundreds of coexisting strains. High rates of turnover endow these communities with a capacity for rapid evolutionary change. Time-resolved sequencing has started to illuminate this process, with several recent studies in mice (1–6) and humans (7–13) documenting genetic variants sweeping through local populations of gut bacteria on timescales of weeks and months. This strain-level variation can alter metabolic phenotypes (2, 14–16), influencing the breakdown of drugs (17) and the invasion of external strains (4, 18). Yet despite their importance, the evolutionary drivers of this *in vivo* adaptation – and their dependence on the host environment – are only starting to be uncovered.

Traditional sequencing approaches have a limited ability to address these questions, since they can only observe the handful of lineages that manage to reach appreciable frequencies within a host. By this time, successful lineages have often acquired multiple distinct mutations (4, 7, 13). This makes it difficult to resolve their underlying fitness benefits, or the pleiotropic tradeoffs that they encounter in different host conditions (2). It also prevents us from observing the other contending mutations that – through a combination of luck and merit – were outcompeted before they were able to reach appreciable frequencies within their host.

Barcoded lineage tracking provides a powerful alternative, enabling quantitative fitness measurements of thousands of independent mutations within a single population (19). However, existing methods for high-throughput isogenic barcoding require specialized genetic tools, and have previously been limited to laboratory strains of yeast (19, 20) and *E. coli* (21). Here we show that similar evolutionary inferences can be obtained from genome-wide transposon insertion sequencing (Tn-Seq) libraries (22–26), which are routinely employed in functional genomics settings. Tn-Seq libraries are traditionally used to identify conditionally essential genes in various bacterial species and environments (22–24, 27), including several recent *in vivo* studies in gnotobiotic mice (16, 25, 28, 29). Here we sought to exploit this same technique as a crude form of barcoding, by focusing on the vast majority of Tn insertions in genes without obvious growth defects. We reasoned that the fine-scale dynamics of these lineages could provide a scalable approach for measuring *in vivo* evolutionary forces in complex communities like the gut microbiota.

## Results

### Fine-scale dynamics of Tn-Seq lineages reveal rapid *in vivo* evolution

To test this hypothesis, we reanalyzed data from a previous transposon screen of multiple commensal gut bacteria in gnotobiotic mice (28). Tn-Seq libraries of four human *Bacteroides* strains – *B. cellulosilyticus* (*Bc*), *B. ovatus* (*Bo*), and two strains of *B. thetaiotaomicron* (*Bt-VPI* and *Bt-7330*) – were combined with 11 other species and gavaged into 20 individually caged mice. Mice were were maintained on either a low-fat/high-plant polysaccharide diet (LF/HPP), a high-fat/high-sugar diet (HF/HS), or alternating sequences of the two (HLH/LHL) for 16 days, with Tn-Seq measurements performed on fecal samples collected at three timepoints (Fig. 1A). Wu *et al* (28) used these data to show that ~10-30% of gene knockouts displayed a consistent fitness cost in at least one of the diets during the first 16 days of colonization. After excluding the Tn insertions in these and other “fitness determinant” genes, we identified a collection of ~60,000-150,000 mutants in each library that were suitable for high-resolution lineage tracking (SI Section 1). By monitoring the relative frequencies of these Tn lineages over time, we sought to quantify the additional evolutionary forces that acted within these populations during the first two weeks of colonization.

**Figure 1:**
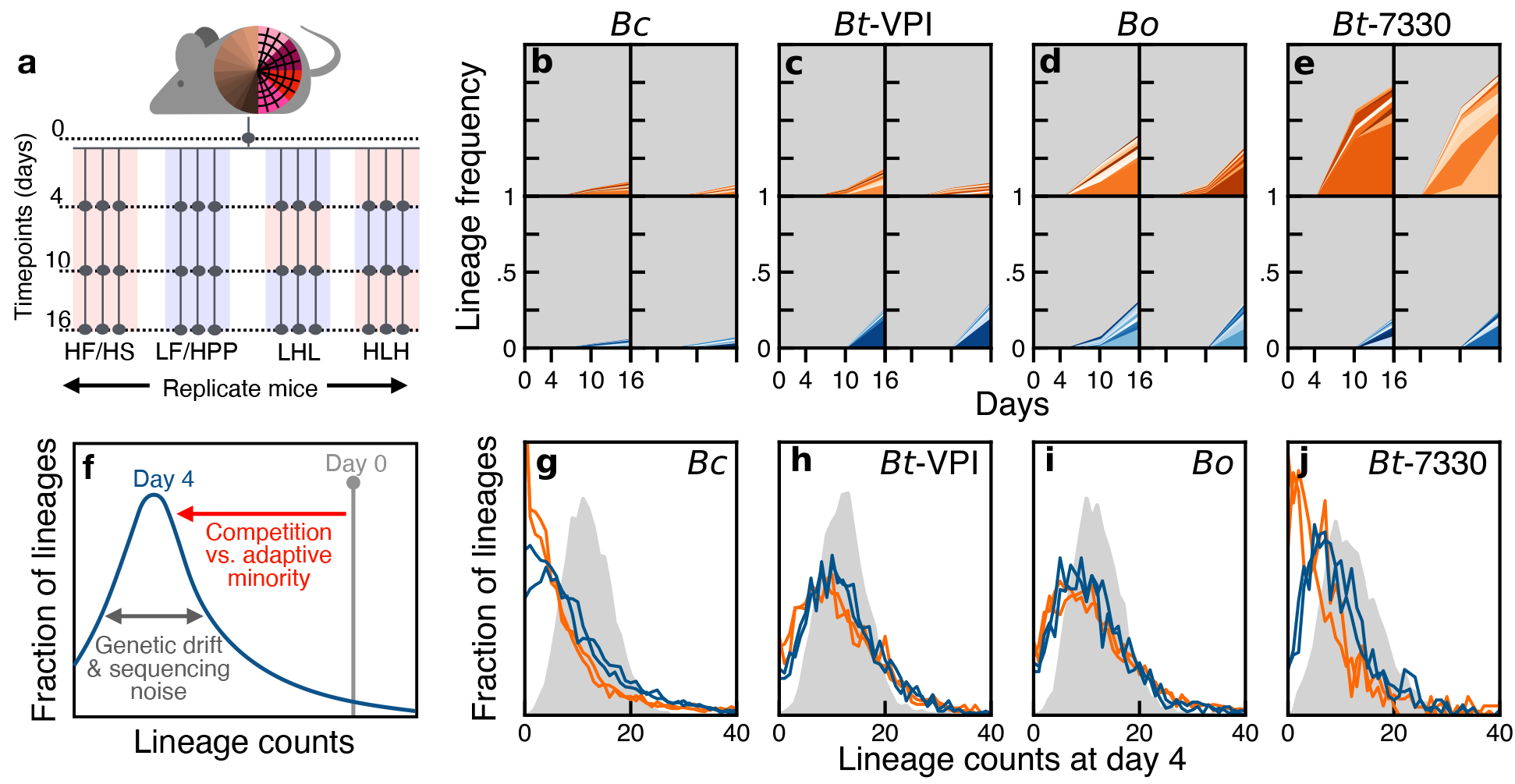
Collective behavior of Tn lineages reveals rapid *in vivo* evolution in gnotobiotic mice. **(a)** Schematic of Tn-Seq experiment in Ref. (28). Mutant libraries of 4 *Bacteroides* strains were introduced into gnotobiotic mice and sequenced over time in different diets. **(b-e)** Individual frequency trajectories of the 10 largest lineages at day 16 in each of two representative mouse from the HF/HS (orange) or LF/HPP (blue) diets. **(f)** Schematic of the simplest evolutionary null model (SI Section 3), where neutral lineages decline due to competition with fitter lineages in the population and stochastic fluctuations from genetic drift and sequencing noise. **(g-j)** Distribution of lineage read counts on day 4 for a subset of the lineages with similar initial frequencies in the input library. Colored lines show the distributions for each of the four mice in (b-e). Grey distributions show the null expectation from sequencing noise alone (SI Section 3).

Consistent with previous observations in other bacterial species (1–4), we found that a handful of lineages expanded to intermediate frequencies (>1%) *in vivo* by day 16 (Fig. 1B-E), indicating rapid positive selection on a subset of the lineages. Most of these lineages expanded in multiple independent mice, but other Tn insertions in the same gene did not (Fig. S1). This suggests that their fitness benefits did not derive from the original Tn insertion, but rather from secondary mutations that accumulated in the library prior to colonization. The consistency across mice contrasted with the variation we observed across the different *Bacteroides* species: the highlighted lineages in Fig. 1B-D accounted for ~50% of the population in *Bt-7330* and *Bo* on day 16, but comprised a much smaller fraction in *Bc* and *Bt-VPI*. This shows that the rates of adaptation in the murine gut can vary between similar gut species, and even between strains of the same species.

These differences between species were much less pronounced on day 4, with no lineage reaching >5% frequency, and most remaining <0.01% (Fig. 1B-D). While low read counts make it difficult to follow these lineages individually, we reasoned that their collective behavior could still encode information about the evolutionary forces operating on these shorter timescales. As a first step, we focused on the subset of lineages that were present at a given frequency *f*_0_ in the initial library, and examined the distribution of their frequencies at day 4 (Fig. 1G-J). In the simplest evolutionary null model (SI Section 3), the typical frequencies of these neutral lineages would decline due to competition with fitter mutations in the population, as well as from stochastic fluctuations from genetic drift and sequencing noise (Fig. 1F).

The observed distributions were largely consistent with this prediction. We found that these aggregated dynamics were remarkably similar across mice in the same diet, and to a lesser degree, between diets as well. However, we once again observed dramatic differences between the various *Bacteroides* species. By day 4, most lineages remained close to their initial frequencies in *Bo* and *Bt-VPI*, while the majority of lineages substantially declined in *Bc* and *Bt-7330*. These differences could not have been caused by genetic bottlenecks, since we found that many of the same lineages were consistently present – and often expanded – in multiple independent mice (Fig. 2A). These correlations indicate that the collective behavior in Fig. 1G-J must have been driven by positive selection within these populations during the first four days of colonization.

**Figure 2:**
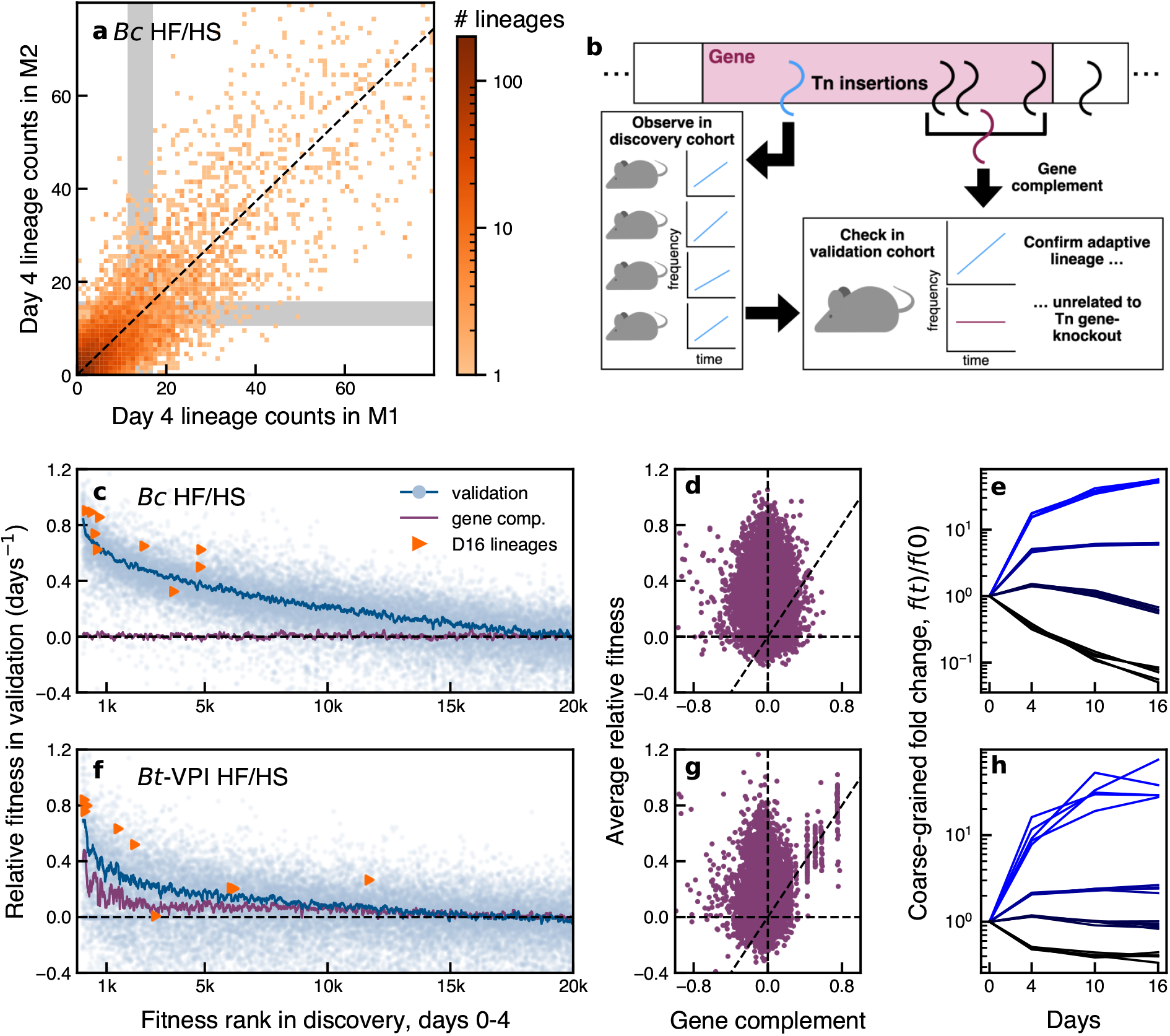
Positive selection on thousands of lineages that are unrelated to their original gene knockouts. (**a**) Joint distribution of day 4 read counts for a subset of *Bc* lineages in two representative mice. Lineages were drawn from a narrow range of initial frequencies in the input library (grey regions). (**b**) Schematic of cross-validation approach for detecting adaptive lineages. (**c,f**) Relative fitness in the validation mice for the fittest 20,000 lineages in the discovery cohort in *Bc* (**c**) and *Bt-VPI* (**f**). Symbols denote individual Tn lineages, while lines denote running averages of 100 lineages (blue) or their corresponding gene complements (purple) (SI Section 4). Triangles indicate the 10 largest lineages in the HF/HS mice on day 16. (**d, g**) Relative fitness of putatively adaptive lineages vs their corresponding gene complement. (**e, h**) Putatively adaptive lineages continue to expand over time. Colored lines show the total frequency of ranks 1-1,000, 1,000-10,000, and 10,000-20,000 (light blue to dark blue), as well as the remaining lineages (black) in different mice.

Intriguingly, the apparent rates of adaptation in Fig. 1G-J were different from those in Fig. 1B-E. For example, *Bc* showed the strongest signatures of positive selection on day 4, but had the smallest number of high-frequency lineages at day 16. *Bo* displayed the opposite trend. This shows that the dynamics of common variants do not necessarily reflect the broader adaptation occurring within these populations at lower frequencies.

### Contrasting the spectrum of adaptive mutations in different gut species

To quantify this adaptive landscape more systematically, we turned to a cross-validation approach that took advantage of the replicated experimental design. We ranked each lineage by its average fold change in a “discovery” cohort of mice on a given diet, and compared this to its relative fitness in a separate “validation” cohort from the same diet (Fig. 2B; SI Section 4). We reasoned that if the expanding lineages were driven by selection on preexisting variants, then their fitness in the validation cohort should be consistently positive as well. Fig. 2C shows an example of this approach for the *Bc* populations in the HF/HS diet between days 0 and 4. While individual lineages were noisy as expected, we nevertheless observed a clear enrichment in positive relative fitness among the top ~15,000 lineages. This suggests that >10% of all lineages in *Bc* experienced strong positive selection during the first four days of colonization (Fig. S2). Moreover, this cohort of putatively adaptive lineages continued to expand over days 4-16 (Fig. 2E), suggesting that their collective fitness benefits were not confined to this initial time interval.

In principle, these *in vivo* fitness benefits could be caused by the gene-knockout effects of the original Tn insertions. Under this hypothesis, we would expect that the other lineages with insertions in the same gene should also expand over the same time interval (Fig. 2B). Surprisingly, however, we observed no strong correlation between the relative fitnesses of the putatively adaptive lineages in *Bc* and the fitnesses of their corresponding “gene complements” (Fig. 2C-D, SI Section 4). This suggests that their *in vivo* fitness benefits were caused by secondary mutations that accumulated in the library prior to colonization. Our analysis shows that the fitness benefits of these mutations are large by evolutionary standards (>10% per day), and are comparable to the “fitness determinants” detected in the original transposon screen (Fig. S3). We also observed considerable variation in fitness within the subset of adaptive lineages (Figs. 2C,E and S4) suggesting that their benefits derived from different underlying mutations.

Similar signatures were present in the other *Bacteroides* species, though the number and magnitudes of the fitness benefits were somewhat different (Figs. 2F, S5, S6). For example, the number of strongly expanding lineages in *Bt-VPI* was lower than in *Bc*. Many of these lineages were also clustered in the same genes (Fig. 2G), suggesting that their fitness benefits were caused by their original Tn insertions. However, even in this case, we found that loss-of-function variants accounted for only a small fraction of the putatively adaptive lineages, since thousands of other lineages expanded by smaller amounts (Fig. S4). *Bo* and *Bt-7330* (Fig. S5) showed similar trends. In each of these cases, we found that the largest lineages at day 16 were enriched among the putatively adaptive mutations at day 4. Interestingly, however, these eventual winners were not necessarily the fittest lineages early on, suggesting that further mutations or environmental shifts were required to reach their dominant frequencies. This highlights how chance and competition among numerous low frequency variants can play an important role in determining which mutations rise to appreciable frequencies within a host.

### Pleiotropic fitness tradeoffs across time and between diets

We next examined how this adaptive landscape varied over time and in different dietary conditions. We observed that the relative fitnesses of individual lineages were remarkably consistent across the HF/HS and LF/HPP diets during the first four days of colonization (Figs. 3A,H-K and S7). This shows that the thousands of adaptive lineages in Fig. 2C,F were not specific to host diet. Intriguingly, the *in vivo* relative fitnesses in *Bc* were also highly correlated with *in vitro* fitnesses measured in several media (Fig. S8), suggesting that they were also not specific to the complex features of their host environment.

**Figure 3:**
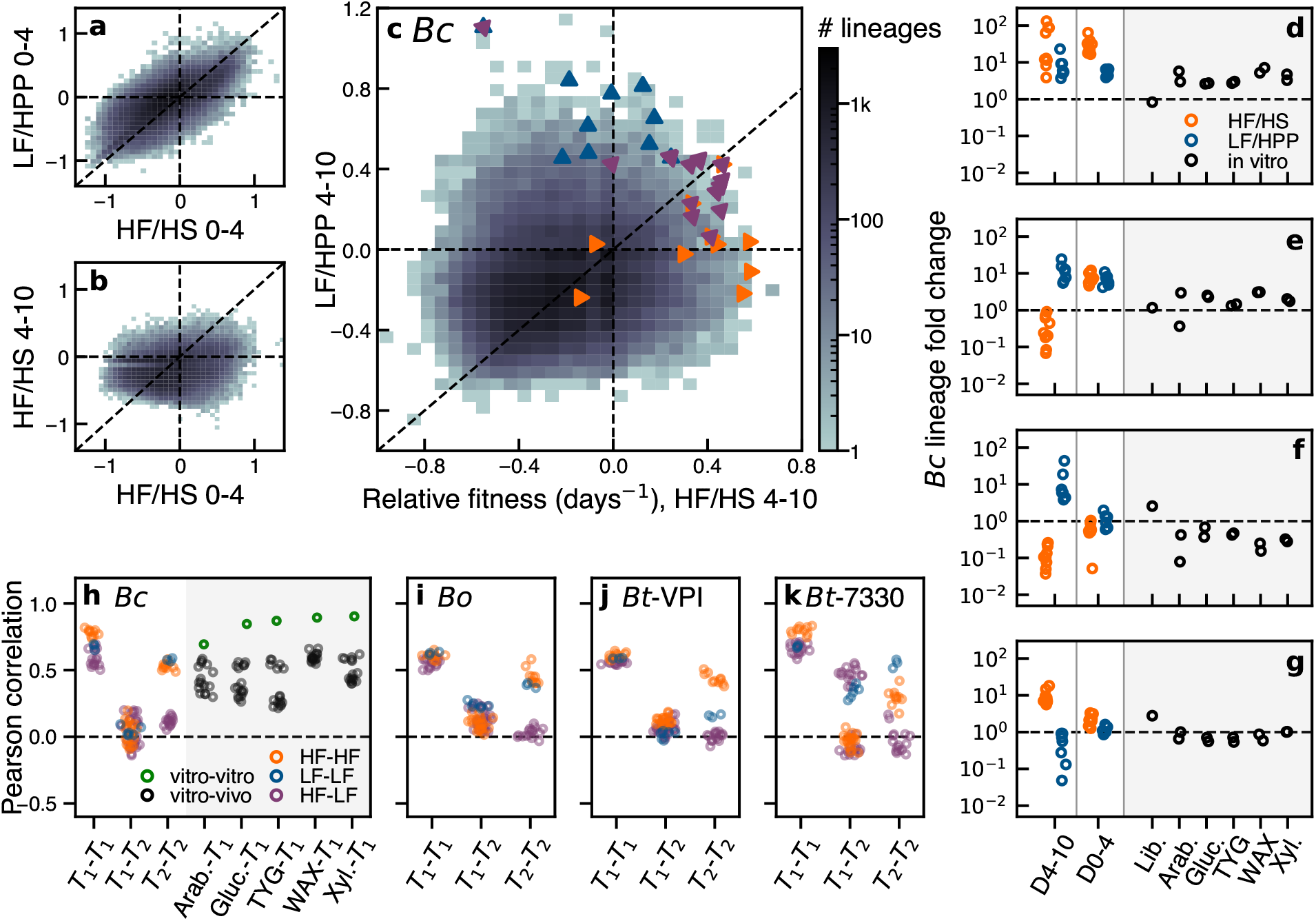
Selection pressures shift over time to reveal diet-dependent fitness tradeoffs. (**a-c**) Joint distribution of relative fitnesses in (**a**) HF/HS vs LF/HPP diets over days 0-4, (**b**) days 0-4 vs 4-10 in the HF/HS diet, and (**c**) HF/HS vs LF/HPP diets over days 4-10; triangles indicate the 10 largest lineages at day 16 in the HF/HS (orange), LF/HPP (blue), or alternating (purple) diets. (**d-g**) Example lineages with strong fitness tradeoffs in different *in vivo* and *in vitro* conditions; circles indicate independent mice or *in vitro* cultures. (**h-k**) Pearson correlation coefficients of relative fitness values of lineages across different pairs of environments. Symbols denote comparisons between individual pairs of replicates. (SI Section 5). *T*_1_=days 0-4; *T*_2_=days 4-10; Lib.=library creation; Arab.=arabinose; Gluc.=glucose; Xyl.=xylose.

In contrast, we found that the relative fitnesses of the lineages were only weakly correlated across time intervals in each of the *Bacteroides* species (Fig. 3B,H-K). This lack of correlation was not driven by the absence of selection at later times: Figs. 3H-K and Fig. S9 show that the relative fitnesses during days 4-10 were still well correlated within the same diet. Instead, these results indicate that the selection pressures shifted over time, potentially reflecting a transition between colonization-dominated vs competition-dominated selection.

Consistent with this hypothesis, we found that the relative fitnesses in later time intervals were only weakly correlated across diets (Figs. 3C,H-K), suggesting that host diet can have a substantial impact on selection at later times. Across the larger set of adaptive lineages, we identified hundreds of individual examples with strong fitness tradeoffs in different dietary conditions (Figs. 3D-F, Fig. S10). Some lineages expanded 10-fold in HF/HS, but were neutral or deleterious in LF/HPP (Fig. 3F); other lineages displayed the opposite trend (Figs. 3D,E). These same lineages exhibited diverse behaviors in other time intervals as well: while the examples in Figs. 3D and E displayed similar tradeoffs between days 4-10, only one of them expanded between days 0-4, while the other was effectively neutral. Conversely, the examples in Figs. 3E and F were both effectively neutral between days 0-4, but exhibited opposing tradeoffs between days 4-10. This diverse range of behaviors provides further evidence that the adaptive lineages were driven by different underlying mutations, which can be differentially amplified by specific sequences of environments.

Despite these strong tradeoffs for individual lineages, our broader characterization revealed no strong evidence for a *global* tradeoff in the underlying fitness landscape. We found that many individual lineages consistently expanded in both diets (Figs. 3G and S10), demonstrating that it is possible for evolution to improve fitness in both environments simultaneously. The lone exception was the comparison with the effective fitness during Tn library generation, which was anti-correlated with *in vivo* fitness in *Bc* and *Bt-7330* (Fig. S11; SI Section 4). These limited anti-correlations suggest that the long-term tradeoffs observed at the population level (2) might not necessarily reflect an underlying physiological constraint, but may actually be an emergent property of their *in vivo* evolutionary dynamics (30).

A striking example of this behavior is illustrated by the handful of lineages that reached the largest frequencies by the end of the experiment. These lineages provide a proxy for the mutations that are likely to dominate the population at long times. We found that the largest lineages in the constant diets exhibited an apparent fitness tradeoff in *Bc*, with higher fitnesses in their home environment and average fitnesses in the other (Fig. 3C). In contrast, the alternating diets consistently selected for lineages that were fitter in both environments, despite their lower overall representation in the underlying fitness distribution (Figs. 3C and S12). This illustrates how clonal competition and fluctuating selection pressures combine to determine the emergent fitness tradeoffs within a population.

## Discussion

Together, these results show how the fine-scale dynamics of genome-wide transposon libraries can enable quantitative inferences of *in vivo* evolutionary forces. We found that the early stages of colonization can be dominated by an intense competition between thousands of adaptive variants – most of which would not be observed with traditional sequencing approaches. While we have observed these dynamics in native human gut strains, it is possible that the high rates of adaptation we observed here could be driven by the novelty of the murine host, or the comparatively low diversity of the artificial gut community. Our approach could be used to test these hypotheses in future experiments, by examining how the spectrum of beneficial mutations differs in communities with higher levels of taxonomic diversity (31).

A key limitation of this approach is that it does not provide direct information about the genetic targets of adaptation. Future experiments could begin to map these molecular drivers by isolating and sequencing a subset of the adaptive lineages we have identified (32). Fig. 3 also suggests that it may be possible to cluster the phenotypic impacts of these mutations directly, by examining their pleiotropic tradeoffs across a large panel of *in vitro* conditions (27, 29). For example, using the existing environments in Fig. 3, we can identify subsets of adaptive mutations whose pleiotropic tradeoffs strongly resemble the gene-level profiles of unrelated genes (Fig. 4A,B; SI Section 6). This suggests that the fitness benefits of these lineages were caused by loss-of-function mutations in the associated genes or pathways, providing a link to their underlying function.

**Figure 4:**
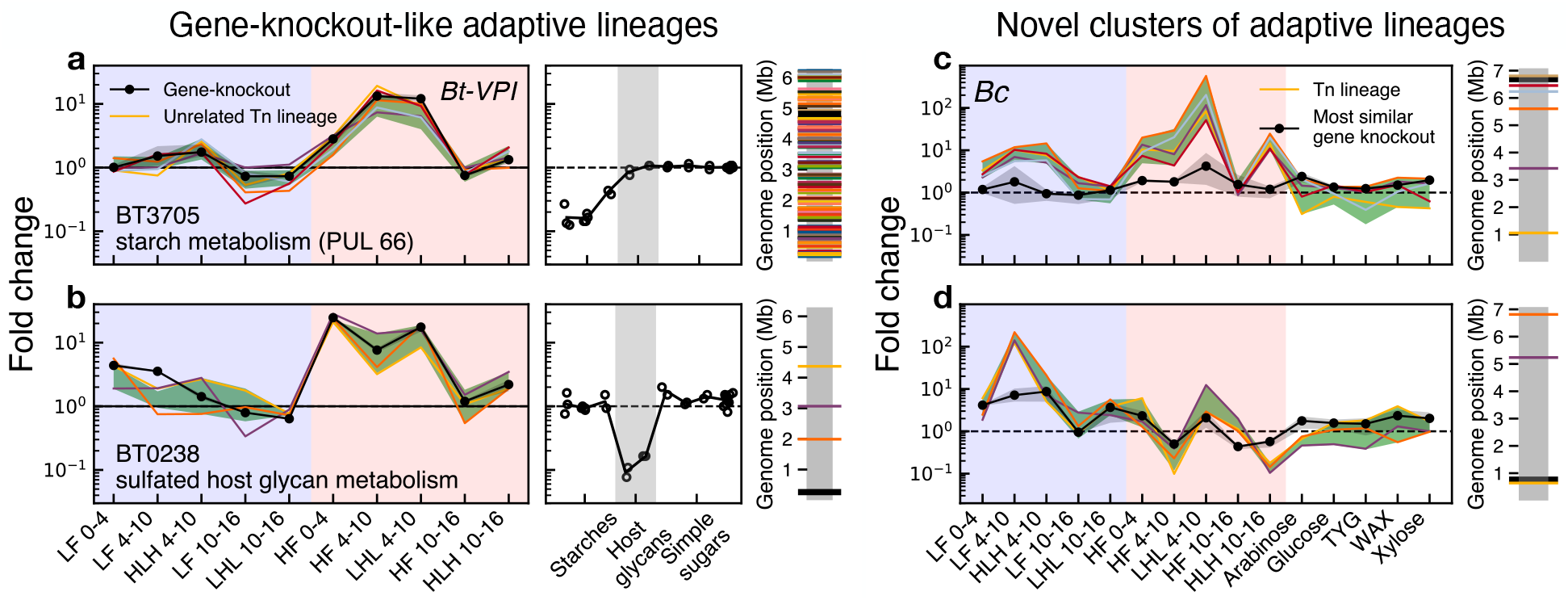
Inferring the functional targets of adaptation using pleiotropic fitness tradeoffs across many environmental conditions. (**a,b**) Examples of adaptive lineages whose fitness profiles closely resemble loss-of-function mutations in other genes. Left panels show the fitness profiles across the entire set of *in vivo* environments in Fig. 3 for a subset of the lineages in the cluster; shaded regions denote an estimated interquartile range of the clustered lineages (green) and the estimated uncertainty of the corresponding gene-level profile (black, SI Section 6). Middle panels show the fold-change of the same gene in several additional *in vitro* monoculture conditions, which were measured in a separate study (29); circles denote independent replicates of the competition assay. Right panels show the location of the target gene (black) relative to the other Tn lineages in the cluster (colors). (**c,d**) Examples of novel phenotypic clusters that do not resemble any gene-level profile. Colored lines are the same as in (**a,b**), while the best-matching gene-level profile (SI Section 6) is shown in black; in both panels, the best-matching gene is >30kb away from any lineage in the cluster.

Using this approach, we identified a cluster of 235 adaptive lineages in *Bt-VPI* that strongly resembled loss-of-function mutations in the polysaccharide utilization locus *PUL66* (Fig. 4A), which encodes genes responsible for starch metabolism (33–35). The disruption of *PUL66* was previously shown to be a fitness determinant in the HF/HS diet in the original Tn-Seq screen (28); Fig. 4A shows that ΔPUL66-like phenotypes are also easily accessible via secondary mutations (though they are still not the fittest lineages in the population). A contrasting example is provided by the sulfatase-maturating enzyme anSME (Fig. 4B), which is important for the utilization of mucin and other sulfated host glycans (36–38). While the disruption of anSME provided significantly larger fitness benefits than Δ*PUL66* in the HF/HS diet (equivalent to an extra ~8-fold increase by day 16), we observed only a handful of adaptive lineages that exhibited fitness profiles similar to ΔanSME (Fig. 4B). The larger number of ΔPUL66-like lineages allowed them to collectively account for a larger fraction of the *Bt-VPI* population at the end of the experiment (~3% vs <1%), despite their smaller initial fitness benefits. This illustrates how differences in mutational accessibility can play a crucial role in determining the phenotypic response of an evolving population.

In addition to these annotatable examples, we also identified recurrent targets of selection whose fitness profiles were distinct from any gene-level knockouts (Fig. 4C,D; SI Section 6). These novel phenotypic clusters were particularly prevalent in *Bc*, which exhibited larger numbers of adaptive lineages (and fewer beneficial gene knockouts) compared to *Bt-VPI*. These different examples in Fig. 4 show how the secondary mutations identified by our approach can illuminate regions of the adaptive landscape that are not accessible via traditional knockout screens.

Finally, while our present analysis has focused on the dominant signal of selection on standing variation, it is possible to extend this approach to identify signatures of *de novo* mutations (Figs. S15 and S16) and rates of genetic drift (Fig. S17). This suggests that future applications of genetic barcoding could be a promising tool for resolving *in vivo* evolutionary forces in complex microbial communities.

## Acknowledgments

We thank M. Wu and J. Gordon for providing access to the original data, and S. Walton, Z. Liu, and other members of the Good lab for useful discussions and feedback on the manuscript. This work was supported in part by the Alfred P. Sloan Foundation grant FG-2021-15708, NIH NIGMS Grant No. R35GM146949, and a Terman Fellowship from Stanford University. B.H.G. is a Chan Zuckerberg Biohub Investigator.

## Author contributions

Conceptualization: D.P.W. and B.H.G.; theory and methods development: D.P.W. and B.H.G.; analysis: D.P.W. and B.H.G.; writing: D.P.W. and B.H.G.

## Data and materials availability

Postprocessed data described in the paper are presented in the supplementary materials. Raw sequencing data are available at the European Nucleotide Archive and NCBI using the accessions provided in the supplementary materials. All analysis code is available on Github (https://github.com/bgoodlab/adaptation_tnseq).

**Supplementary Materials**

Supplemental Methods

Table S1

Fig S1 – S15

## Supplemental Methods

### 1 Data used in analysis

The raw data used in this work were obtained from a previous study (28), in which libraries of 4 human Bacteroides strains (*Bc*, *Bo*, *Bt-VPI*, and *Bt-7330*) were combined with 11 other species and gavaged into gnotobiotic mice. Multi-taxon transposon insertion sequencing (InSeq) was performed on each input library (with 23-41 technical replicates per species), as well as on fecal samples taken on days 4, 10, and 16. One of the species (*Bc*) was also assayed in a variety of *in vitro* conditions. The full list of samples and technical replicates is provided in Table S1.

Raw sequencing reads from the InSeq experiments were downloaded from the European Nucleotide Archive (accession no. PRJEB9434), and the raw reads from the input libraries were downloaded from https://gordonlab.wustl.edu/INSeq_input_sample_reads/. Reference genomes for each of the 4 *Bacteroides* strains were obtained from National Center for Biotechnology Information (accession no. PRJNA289334). After removing the transposon sequence, each read was matched to its corresponding insertion location on the reference genome using using a custom Python script. Only exact matches were retained, and reads that matched to multiple locations were excluded. We assumed that each unique location *ℓ* corresponded to a distinct lineage founded by a single transposon insertion event, and we calculated the total number of reads *R_ℓ,s_* corresponding to lineage *ℓ* in sample *s*. These read counts formed the basis of all of our downstream analysis.

For each *Bacteroides* strain, Wu *et al*. (28) determined a set of “fitness determinant” genes whose Tn-insertion knockouts were estimated to be non-neutral. Most of these gene knockouts had deleterious fitness effects, though a small fraction were beneficial [see Tables S4A-B, S9A-B, S14A-D of Ref. (28)]. To arrive at a conservative set of quasi-neutral Tn lineages, we removed from downstream analysis all lineages whose corresponding transposons fell within or 100bp upstream of any of the fitness determinant genes identified by Wu *et al* (28). This filtering step left a total of *n*=418,879 lineages across the four libraries (88,396 in *Bc*; 117,020 in *Bo*; 150,849 in *Bt-VPI*; and 62,614 in *Bt-7330*), which we used for all of our subsequent analysis. We estimated the relative frequencies of these remaining lineages using the plug-in estimator,

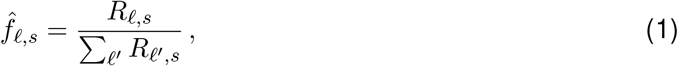

where the denominator sums over all of the filtered lineages within a given *Bacteroides* strain. This renormalization scheme ensures that the relative fitnesses inferred in our later analyses are independent of the fitness determinant genes examined in Ref. (28). (The sole exceptions are Fig. 4 and Figs. S3,13, and 14, which compare the relative fitnesses of the fitness determinant genes identified by Wu *et al*. (28) with the additional adaptive lineages identified in the present work.)

### 2 Evolutionary model of lineage dynamics

We assumed that the temporal dynamics of the Tn lineages could be described by a simple evolutionary model, in which the lineages within a given mouse *m* competed with each other as a well-mixed population. In the most general form of this model, the frequencies of rare lineages (*f_ℓ,m_* ≪ 1) are described by a system of coupled stochastic differential equations,

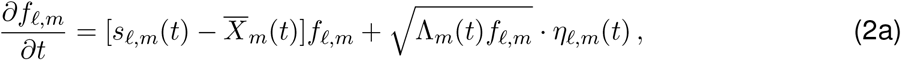

where *η_ℓ,m_*(*t*) is a Brownian noise term with mean zero and variance one (39), and Λ_*m*_(*t*) is the strength of genetic drift in mouse *m* at time *t*. Each lineage *ℓ* has instantaneous fitness *s_ℓ,m_*(*t*), while 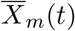 is the mean fitness of the population,

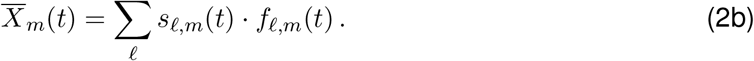

Equation (2) is a time-dependent generalization of the branching process model employed in previous *in vitro* lineage tracking studies (19, 20). The additional time-dependence allows us to account for shifting selection pressures and population bottlenecks that might arise in more complex *in vivo* settings.

If the functions *N_m_*(*t*), *s_ℓ,m_*(*t*), and 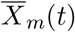 are known, the dynamics of an individual lineage in Eq. (2) can be solved using standard techniques (20). Given an initial frequency *f_ℓ,m_*(*t_0_*) at time *t_0_*, the moment generating function for the lineage frequency at a later time *t* is given by

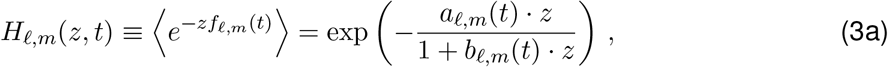

where the functions *a_ℓ,m_*(*t*) and *b_ℓ,m_*(*t*) are defined by

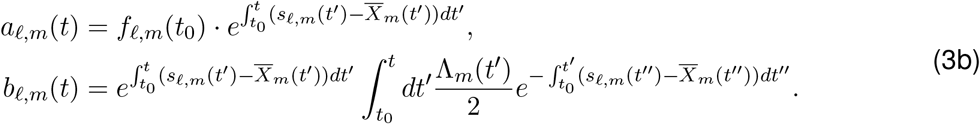

The mean and variance of *f_ℓ,m_*(*t*) can then be obtained from the derivatives of *H_ℓ,m_*(*z,t*):

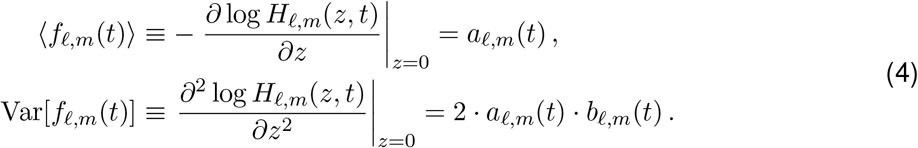

We assumed that the observed read counts *R_ℓ,s_* were generated from these lineage frequencies through an additional sampling process, which encapsulates the combined effects of cell sampling, PCR amplification, and DNA sequencing. We assumed that this sampling process is unbiased, so that on average,

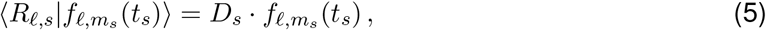

where *D_s_* is the total coverage of sample *s*. Similarly, the variance can be written in the general form,

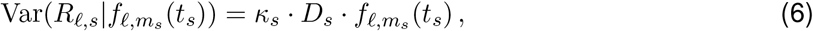

where *κ_s_* is a constant that describes the deviations from simple Poisson sampling. While the full distribution of the sampling process can be complicated, previous work (19, 20) has shown that it can often be approximated by a second branching process with a conditional generating function

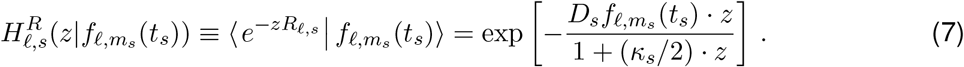

By marginalizing over the random value of *f_ℓ,m_s__*(*t_s_*) using Eq. (3), we can obtain a corresponding expression for the marginal distribution of the read counts

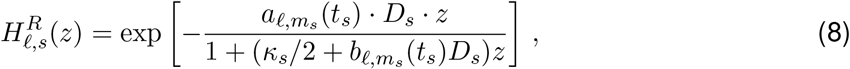

whose mean and variance are given by

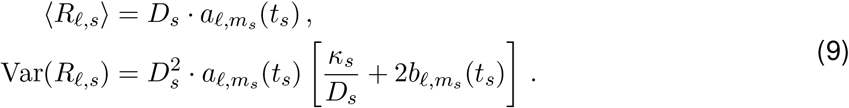

In this way, Eq. (8) provides a model that links the observed read counts *R_ℓ,s_* to the underlying evolutionary parameters *N_m_*(*t*), *λ_m_*(*t*), *s_ℓ,m_*(*t*), and 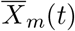 in each mouse. All of our subsequent analyses were derived by considering different limits of this basic model.

### 3 Distributions of lineage frequency shifts

Our initial expectations for the distribution of lineage frequency shifts in Fig. 1G-J were informed by the simplest limit of Eq. (2), in which the vast majority of the focal lineages are effectively neutral (*s_ℓ,m_*(*t*) ≈ 0). For lineages with similar initial frequencies (*f_ℓ,m_*(*t*_0_) ≈ *f*_0_), the branching process parameters in Eq. (3) reduce to a pair of lineage-independent functions,

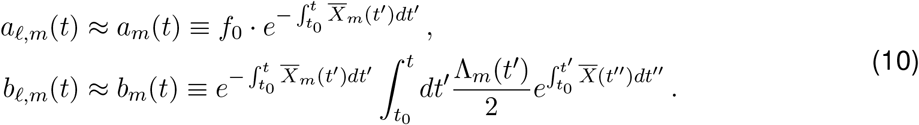

Note that the mean fitness 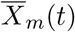 can still be non-zero if a small minority of the lineages in the population have non-zero fitness. In this way, the statistical behavior of a large number of neutral marker lineages can in principle provide information about the population parameters 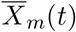 and *N_m_*(*t*) (19).

For example, by substituting Eq. (10) into Eq. (8), we see that the average size of a neutral marker lineage declines over time due to competition with fitter lineages in the population 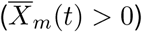. Likewise, the variance in the observed read counts grows due to a combination of genetic drift (∝ 1/*N_m_*(*t*)) and sequencing noise (∝ *κ_s_*/*D_s_*). However, it can be difficult to apply these heuristics in practice, since the mean and variance can be biased if a small number of highly fit lineages happen to be present in the initial pool. We therefore turned to other characteristics of the lineage frequency distribution that are more robust to small amounts of “contamination” by non-neutral lineages.

For example, in the limit that the lineage fluctuations are small (Var(*R_ℓ,s_*) ≲ *D_s_f*_0_), one can invert Eq. (8) to obtain an asympotic expression for the probability distribution of *R_ℓ,s_* (19),

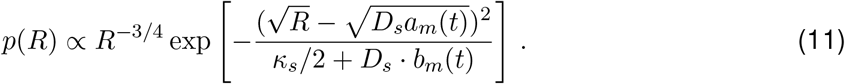

The peak (or mode) of this distribution occurs at a characteristic value *R*\*, which is given by

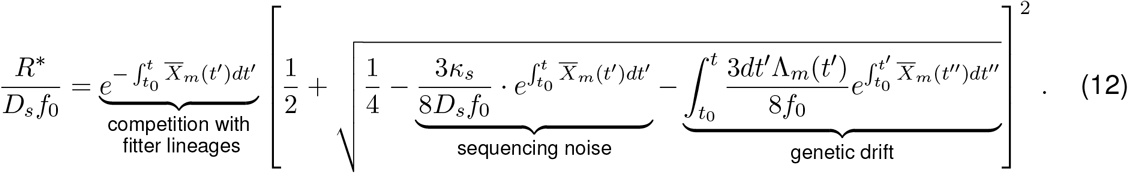

This expression shows that peak of *p*(*R*) will decline due to competition with fitter lineages in the population, as well as through genetic drift and sequencing noise. Similarly, Eq. (11) shows that the characteristic width around this peak will spread out due to genetic drift and sequencing noise. In contrast to the mean and variance above, we expect that these “typical measures” will be robust to the inclusion of a small number of non-neutral lineages in the initial pool.

To compare these predictions with the data, we tabulated the empirical distributions of day 4 read counts for the subset of lineages in each mouse whose day 0 frequencies fell in the range 10/*D*_*m*,4_ ≤ *f*_0_ ≤ 15/*D*_*m*,4_ (Fig. 1G-J). These day 0 frequencies were estimated by pooling all but one of the technical replicates of the input library. As a comparison, we performed the same procedure on the remaining input replicate to obtain an empirical null distribution showing the effects of technical noise alone (Fig. 1G-J). Deviations from this null distribution suggest that the observed dynamics are driven by the evolutionary forces of natural selection and/or genetic drift.

The joint distribution in Fig. 2A was computed using a similar procedure. We identified a subset of lineages with similar initial frequencies, 2 · 10^-5^ < *f*_0_ < 3 · 10^−5^ in the input library, and measured their day 4 read counts in a pair of mice in the same diet. Under our simple model in Eq. (10), this joint distribution should factor into a product of the two marginal distributions [*p*(*R*_1_, *R*_2_) ≈ *p*(*R*_1_)*p*(*R*_2_)], regardless of their individual locations or widths. The strong correlations in Fig. 2A indicate departures from this simple model, in which a substantial fraction of the focal lineages have non-zero fitnesses that are shared across independent mice.

### 4 Cross-validation approach for inferring lineage fitnesses

The strong correlations in Fig. 2A suggested an alternative model, in which a substantial fraction of the lineages have non-neutral fitnesses [*s_ℓ,m_*(*t*) ≠ 0], which are similar for different mice in the same diet [*s_ℓ,m_*(*t*) ≈ *s_ℓ,em_*(*t*)]. If the mean fitnesses are also similar for populations in the same environment 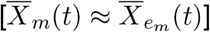, then our expression for the average lineage size in Eq. (9) reduces to

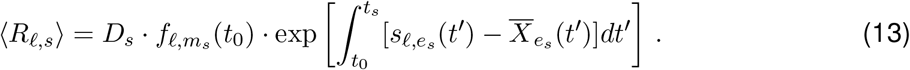

This suggests that it should be possible to infer the underlying fitness functions *s_ℓ,e_*(*t*) by leveraging the independent observations of *R_ℓ,s_* in replicate mice. Due to the difficulty in identifying a set of truly neutral lineages, we chose to work directly with the *relative fitness* 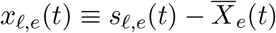, which measures the instantaneous growth rate of a lineage as it competes with its surrounding population. We also defined a time-averaged version of the relative fitness over a given time interval *t*_0_ ≤ *t* ≤ *t*_1_:

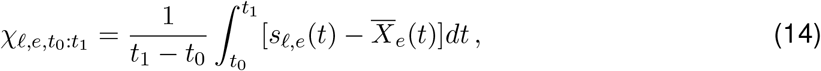

which contains all the information necessary to predict the average fold change of the lineage over that time interval:

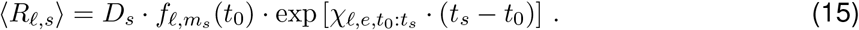

In this way, the general fitness inference problem can be reduced to inferring a finite collection of *χ*_*ℓ,e,t*_0_:*t*_1__ values for each time interval in the experiment. We note that in this notation, temporal variation in *x_∓_*(*t*) and *χ*_*ℓe,t*_0_:*t*_1__ can arise both through changes in the underlying selection pressures *s_ℓe_*(*t*), as well as from the ordinary evolutionary dynamics of the mean fitness 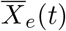. We discuss ways to distinguish between these sources of temporal variability in more detail below.

We estimated the relative fitness *χ*_*ℓe,t*_0_:*t*_1__ by pooling observations from multiple replicate mice in the same environment. For a given cohort of mice 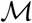, we estimated the relative fitness using the plug-in estimator,

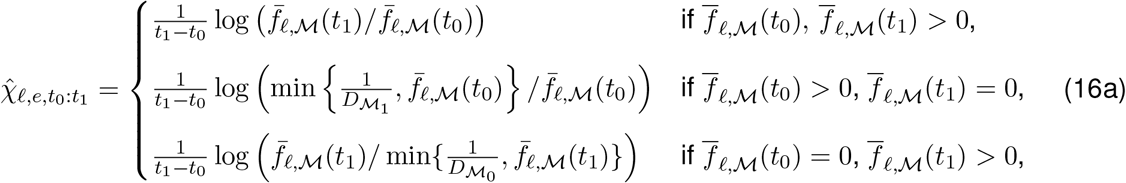

where 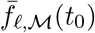 is the weighted average of the lineage’s frequency within the cohort,

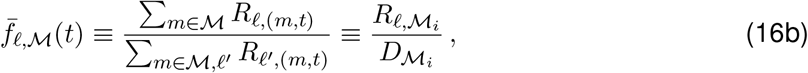

and 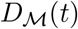 is the total coverage of the library across the cohort of mice,

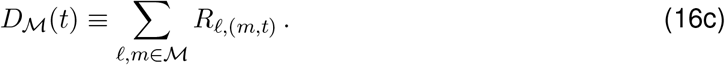

The edge cases in Eq. (16) allow us to assign finite relative fitnesses to lineages with zero reads at either the initial or final time point. In these cases, the min{·} terms act like an effective pseudocount, which is conservatively biased to assign zero relative fitness to lineages with sufficiently low frequency [e.g. *f*(*t*_0_) < 1/*D*(*t*_1_)].

#### *In vitro* fitness estimates

We used a similar approach to estimate relative fitnesses in the *in vitro* environments (Fig. 3H-K and Fig. S8). Wu *et al*. (28) competed the *Bc* library in 5 *in vitro* growth media; 2 independent cultures were inoculated in each medium, and 3 aliquots were sequenced from each culture after reaching stationary phase. We estimated the relative fitness in each independent culture using Eq. (16); since no explicit time interval was reported, we set *t*_1_ – *t*_0_ = 1 to reflect the duration of a typical overnight culture.

We also defined a special *in vitro* environment representing the library creation process. We assumed that all of the Tn lineages were initially founded by unique insertion events in a single cell, so that their frequencies in the input library reflect the differential growth that occurred during the library creation process. We estimated the relative fitness in this effective environment using Eq. (16) with a uniform initial frequency,

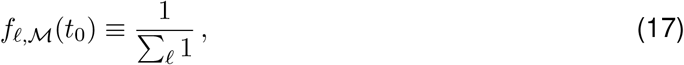

and an arbitrary time interval *t*_1_ – *t*_0_ = 1.

#### Cross validation of fitness estimates

To verify that putatively adaptive lineages measured in a cohort were not statistical outliers due to biological or technical sources of noise, we adopted a cross-validation approach. In this approach, *k* mice that were maintained in same dietary environment *e* were evenly divided into discovery 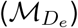 and validation 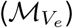 cohorts. The replicate measurements of the input library were also evenly divided between the two cohorts. We used this partitioning to obtain independent estimates of the relative fitness of each lineage in both the discovery and validation cohorts. We ranked each lineage by its fitness in the discovery cohort, and examined how the fitnesses in the validation cohort varied as a function of their rank *ρ*(*ℓ*) in the discovery cohort (Figs. 2C,E and S5). Since the two cohorts have independent sources of technical noise, systematic correlations between these two quantities can be used to distinguish genuine fitness differences from statistical fluctuations in read counts.

To increase the signal-to-noise ratio, we restricted our attention to lineages with initial frequencies >10^−6.5^ (equivalent to ~10 reads in the pooled input library in each species). In addition, we only examined lineages with a minimum number of expected reads in the validation cohort:

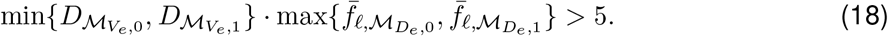

These filters were used to generate the rank-ordered fitness distributions in Figs. 2C,E and S5–S7.

To distinguish systematic trends from the noisy estimates of individual lineages, we coarse-grained groups of lineages based on their relative fitness rank, *ρ*(*ℓ*), in the discovery cohort. For a given range of ranks *ρ*_1_ ≤ *ρ* ≤ *ρ*_2_, we defined the coarse-grained frequency 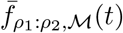 by summing over the individual lineage frequencies in Eq. (16b):

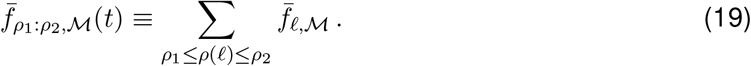

Under the simple model in Eq. (15), this coarse-grained frequency will grow as

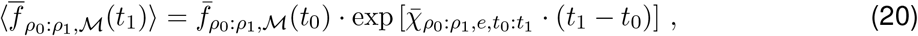

where 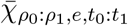 is the average relative fitness,

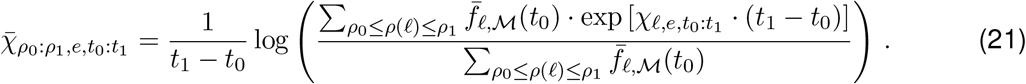

Positive values of 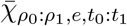 indicate that at least some lineages in the coarse-grained grouping have positive relative fitness. We estimated this coarse-grained relative fitness in the validation cohort using an analogous version of Eq. (16), in which 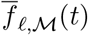 is replaced by 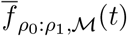. For example, the blue lines in Figs. 2C,E and S5–S7 show the coarse-gained relative fitnesses in the validation cohort in sliding windows of 100 consecutive ranks. The consistently positive values observed at lower values of *ρ*(*ℓ*) indicate that many of the underlying lineages had positive relative fitness.

#### Distinguishing gene-level and lineage-level fitness effects

We used a similar coarse-graining procedure to estimate the relative fitness of the gene complement of each lineage (Fig. 2B). The gene complement was defined for a focal lineage *ℓ* if its transposon insertion fell in the coding region or <100bp upstream of an annotated gene. In this case, the gene complement 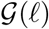 was defined to be the collection of all other lineages (excluding the focal lineage) that also fell within the same gene as *ℓ*. We used this collection of lineages to define a coarse-grained frequency,

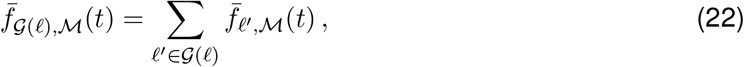

which represents the total frequency of all of the other lineages associated with the same gene. We reasoned that if the relative fitness of lineage *ℓ* was caused by the gene knockout effect of the original Tn insertion, then the dynamics of the gene complement 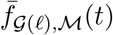 should be statistically similar to the dynamics of the focal lineage 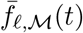. We tested this hypothesis by estimating the relative fitness of the gene complement of each lineage using an analogous version of Eq. (16), with 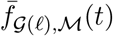 replacing 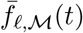. The purple lines in Figs. 2C,E and S5 show sliding averages of the gene complement fitnesses in the validation cohort (calculated using the same coarse-graining scheme described above) as a function of the relative fitness rank of the focal lineage in the discovery cohort. The large separation between the purple and blue lines in these panels indicates that the fitness advantages of many lineages were not caused by their original Tn insertion.

#### Verifying the cross-validation procedure using simulated data

To confirm that our filtering and cross-validation procedure could consistently estimate lineage fitnesses in a population, we simulated lineage dynamics in “biological replicates” designed to mimic the conditions of our experiments. First, we generated an empirical distribution of lineage fitnesses during days 0-4 by averaging across the 9 mice fed the HF/HS diet in this interval in the dataset, as described in SI Section 3. These inferred fitnesses, along with the input frequencies, were used to initialize 9 identical populations. We then simulated these populations for 40 generations (~4 days) under a Wright-Fisher model with a fixed population size *N_e_* = 10^8^. Finally, to generate simulated sequencing libraries, we modeled the sampling and sequencing of each population as a sequence of two Poisson sampling steps, each equal to the empirical sequencing depth at day 4 (which varied across *Bacteroides* species and mice). We then performed the filtering and cross-validation procedures described above to produce rank-order curves from these simulated populations (Fig. S2). These simulations confirmed that our cross-validation approach could reliably distinguish the presence or absence of fitness variation in a population.

#### Estimating the overall number of adaptive lineages

Our simulations showed that the coarse-grained rank order curves reliably estimated the ground truth rank order curves in each of the scenarios we considered. This suggests that these coarse-grained curves can be inverted to obtain an estimator for the true distribution of relative fitnesses. The results are shown in Figs. S2 and S6 for both simulations (Fig. S2) and the observed data (Fig. S6). For visual clarity, the computed distributions were smoothed with Gaussian kernel density estimation before plotting. The resulting distributions provide a direct estimate of the relative numbers of lineages with different relative fitnesses.

The interpretation of these distributions is complicated by the fact that we also observed a large degree of negative fitness variation from day 0-4 (Fig. 3A and Figs. S6–S8), suggesting that there is substantial maladaptive heritable variation in each *Bacteroides* population as well. This makes it difficult to identify the relative fitness that corresponds to the neutral ancestor strain. To be conservative, we therefore only considered a mutation to be adaptive if it increased in frequency over the relevant time interval. This ensures that the lineage is at least as fit as the average fitness of the population.

As an alternative to this approach, we sought to make an independent estimate of the number of adaptive lineages that did not rely on any coarse-graining scheme. First, among the highest *N*_+_ ranked lineages in the discovery cohort, we counted the number of lineages 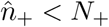 with consistent (positive) relative fitnesses in the validation cohort. We then compared this to a null expectation for the number of lineages that we would have positively cross-validated by chance, i.e. in the absence of correlations across mice. Under the null hypothesis that lineage fitnesses were uncorrelated across discovery and validation cohorts, we drew *N*_+_ lineages randomly among all ranked lineages, and determined the number *n*_+,0_ < *N*_+_ with positive fitnesses in the validation cohort. The number of lineages with positive relative fitnesses observed in excess over the null model,

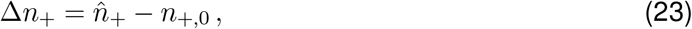

represents a lower bound on the number of truly adaptive lineages among the first *N*_+_ ranks. To see that this is a lower bound, we define the distribution of true relative fitnesses (in the validation cohort) 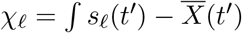 in the (ranked) set of lineages as *ρ*(*χ*), and the probability of measuring a positive relative fitness 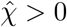 in the validation cohort conditioned on *χ* as 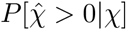. Then, the null expectation 〈*n*_+_, 0〉 from *N*_+_ draws, 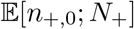, can be expressed as

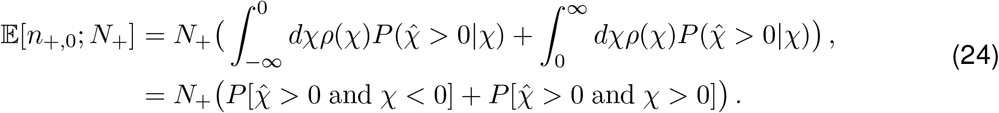

While the empirical null thus discounts from *n*_+_ the expected number of false positives (the first term), it also discounts the expected number of true positives (the second term). Among HF/HS mice, we estimated at least Δ*n*_+_ = 9115 ± 48 adaptive lineages in excess of the null expectation among the first 15,000 ranks in *Bc*, 4074±42 out of 10,000 in *Bo*, 2822±45 out of 10,000 in *Bt-VPI*, and 1630±25 out of 4,000 in *Bt-7330*. Similar estimates were obtained for the LF/HPP diet.

### 5 Inferring joint fitnesses and tradeoffs across environments

To estimate the joint distributions of relative fitnesses across a pair of environments and/or timepoints (Figs. 3A-C and S9), we began by splitting the samples into non-overlapping cohorts for each of the two environments, *e*_1_ and *e*_2_. This division ensured that statistical fluctuations in one environment did not influence relative fitness estimates in the other environment. We used Eq. (16) to estimate the relative fitness in each environment for all lineages that satisfied Eq. (18) in at least one of the two environments.

The pairwise correlations in Fig. 3H-K were estimated from a single biological replicate in each environment, and Pearson correlations were computed for different pairs of biological replicates. However, to avoid biases from our plug-in estimator, in each pair we calculated the correlation only among lineages measured at non-zero frequency in both time points in each replicate.

We defined a lineage as exhibiting a fitness tradeoff if its relative fitness (*χℓ;ℓ,t*_0_:*t*_1_) had opposite signs in a pair of environments or time intervals. To robustly detect such lineages, we took a similar cross-validation approach as described in Section 4 above. For example, to detect the fitness tradeoffs between diets over days 4-10, we first used Eq. (16) to estimate 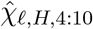 and 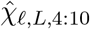 in discovery cohorts of HF/HS and LF/HPP mice. We used these fitness estimates in the discovery cohort to classify each lineage into one of the four quadrants in the (*χ*_*ℓ,H*,4:10_, *χ*_*ℓL*,4:10_) plane: the (+,−) and (−,+) quadrants indicate a potential fitness tradeoff, while the (+,+) quadrant indicates a consistent fitness benefit in both environments. We also defined a quantitative measure of the tradeoff magnitude,

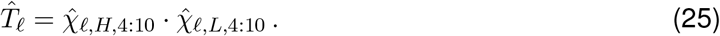

Strong fitness tradeoffs correspond to large, negative values of 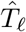, while positive values of 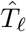 indicate consistent fitness benefits. Several example lineages from *Bc* with large values of 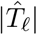 are illustrated in Fig. 3D-F.

To assess the significance of these tradeoffs, we used the validation cohort to check which lineages remained in the same quadrant as their discovery cohort (Fig. S10). To control for multiple hypothesis testing, we then compared the observed number of lineages found in the same quadrant to the following null expectation. If we designated *N_L_* lineages as putatively fit in *L* and unfit in *H* based on their quadrant in the discovery cohort, we drew *N_L_* null lineages without replacement among those with positive *L* fitness in the discovery cohort. We then determined how many of these *N_L_* null lineages were found in the consistent quadrant in the validation cohort. This null model preserves the correlations in positive fitness across mice fed one diet, and assumed that measured tradeoffs in the other diet were simply due to biological or technical noise. We performed an analogous comparison to validate lineages that were putatively fit in *H* and unfit in *L*, drawing null lineages with positive H fitness in the discovery cohort. The excess of lineages with consistent behavior across validation and cohorts over the null expectation in both categories indicated a statistical enrichment for genuine fitness tradeoffs in hundreds of lineages (Fig. S10). We used an analogous approach to identify “generalist” lineages with consistent fitness benefits in both environments, and observed a similar enrichment of these mutations as well (Fig. S10).

### 6 Identifying clusters of phenotypically similar lineages

A key limitation of lineage tracking methods is that they do not provide direct information about the genetic targets of adaptation. This limitation can in principle be overcome by performing whole-genome sequencing on isolated adaptive lineages (4, 32, 40). However, isolating and sequencing the required number of lineages can be challenging, particularly in multi-species settings like Fig. 1, where many adaptive lineages can still reside at low frequencies. Understanding the functional impact of these mutations can also be challenging, since they must be further grouped into larger functional units (32, 41). Here we propose a different approach for annotating mutations that bypasses the intermediate sequencing step, which makes it particularly well-suited for analyzing large numbers of adaptive lineages. This approach is motivated by recent work in functional genomics (27) and experimental evolution (42, 43) showing that the function of genomic variants can often be inferred by examining their pleiotropic tradeoffs across large numbers of *in vitro* environments.

The premise of this approach is that two functionally distinct mutations can have similar fitness effects in one environment, but their fitness effects in other environments may expose differences in their underlying molecular phenotypes. An example of this behavior is shown in Fig. 3: the lineages in panels E and F had similar relative fitnesses in both the HF/HS and LF/HPP diets between days 4-10, but their relative fitnesses diverged between days 0-4. These consistent differences allow us to conclude that the adaptive lineages were likely driven by different underlying mutations.

By the same logic, if two adaptive lineages exhibit the same growth rate differences across a sufficiently large number of environmental conditions, then it is likely that they have acquired either the same mutation or a pair of functionally similar variants (e.g. two mutations in the same gene, or perturbations of a common pathway or module). If the genetic basis of one of these lineages can be determined (e.g. via whole-genome sequencing), then this information can serve as a “functional annotation” for the other lineages in the same phenotypic cluster. Even in the absence of additional sequencing information, the presence of phenotypically similar lineages can provide important information about the degree of evolutionary parallelism within a population (44), and the number of independent phenotypes that can be tuned by natural selection (43). The resulting clusters can also be annotated directly based on the environments in which they exhibit strong fitness effects. Previous work has used this approach to infer the function of poorly annotated genes in diverse species of bacteria (27, 29), by comparing the “fitness profiles” of gene knockouts obtained from genome-wide Tn-Seq screens across a diverse range of environments. Here we show how extensions of this approach can be used to infer the functional targets of adaptation for some of the secondary mutations in Figs. 1–3.

#### Identifying secondary mutations similar to beneficial gene knockouts

As a simple proof of principle, we first searched for adaptive lineages whose fitness profiles resembled one of the loss-of-function variants from the original TnSeq screen (28). We identified these “knockout-like” lineages by comparing the relative fitness profiles of individual Tn lineages (measured across the panel of *in vivo* environments defined below) against the gene-level profiles of unrelated genes. Beneficial lineages whose fitness profiles closely matched one of the gene-level profiles (and had no gene-level signal of their own) were classified as knockout-like lineages, whose secondary mutations might have occurred within the associated gene (or within its associated operon or molecular pathway). We restricted this analysis to *Bt-VPI*, since it exhibited the cleanest examples of beneficial gene knockouts across the 4 *Bacteroides* libraries (data not shown).

To define our set of environments, we took advantage of the fact that the *in vivo* selection pressures varied over time and across host diets (Fig. 3). We therefore split the data into *N_env_* = 10 *in vivo* environments, each of which was a combination of a time interval (Days 0-4, 4-10, or 10-16) and a particular host-diet history (LF/HPP, HF/HS, LHL, HLH). Since LF/HPP and LHL shared the same host-diet history up to day 4 (and likewise HF/HS and HLH), we treated these cases as the same environment between days 0-4.

We defined the coarse-grained fitness profile of a set of lineages 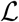 as the vector of log-fold-changes in each environment,

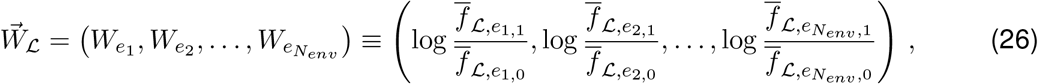

where 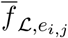. represents the *j*th timepoint of the *i*th environment in a *particular* cohort of replicates 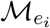, such that

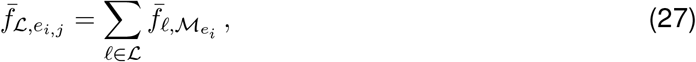

where 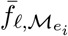 is defined in Eq. (16b). Except where noted, fitness profiles were measured using every replicate available in each environment. Because we are interested in the fitness profiles of beneficial gene knockouts, we measured these frequencies with respect to the whole library, including the fitness determinant genes identified by Wu *et al*. (28); this differs from our analysis in the rest of the study, which excluded lineages that fell in these genes (SI Section 1). We measured the similarity between two fitness profiles using the Euclidean distance,

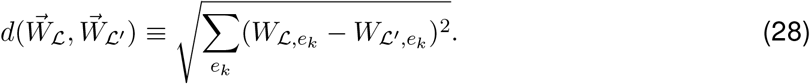

Other metrics, e.g. correlation coefficients, did not qualitatively impact the results.

Next, we curated a reference list of strongly beneficial gene knockouts. We measured the coarse-grained fitness profile 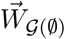 of every gene 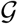 using the Tn lineages falling in 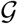 [see also Eq. (22) in SI Section 4]. The notation 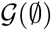 indicates that, unlike the gene complement in Eq. (22), no Tn lineages were excluded from this coarse-grained fitness profile. Based on these results, we restricted our attention to the 88 gene knockouts that exhibited a fold change >10 in at least one environment (i.e., 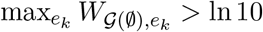), or a fold change >5 in at least two environments. Our earlier analysis suggested that a substantial fraction of these putatively beneficial gene knockouts could be driven by the expansion of a Tn lineage that had acquired an adaptive secondary mutation. Hence, for each of the remaining gene knockouts, we re-calculated a set of “leave-one-out” fitness profiles, 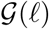, that excluded each lineage *ℓ*:

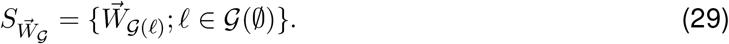

We reasoned that if a single adaptive lineage drove the coarse-grained fitness of a gene, then excluding this lineage would dramatically reduce the leave-one-out fitness profile of the gene. Conversely, if the gene’s coarse-grained fitness profile was driven by many similar lineages, then leaving out any one lineage should not dramatically change the knockout fitness profile. Based on this logic, we removed from consideration any gene which contained a leave-one-out fitness profile that did not exhibit a fold-change >5 in one environment or a fold-change >3 in two environments. This procedure eliminated a majority of the remaining genes; visual inspection of leave-one-out fitness profiles (or individual lineage trajectories) confirmed that nearly all were driven entirely by the behavior of single Tn lineages. This left a tractable list of 32 high-quality beneficial gene knockouts in *Bt-VPI*, many of which were flagged as beneficial fitness determinant genes by Wu *et al*. (28).

We used an empirical approach to determine the relevant distance threshold (*d*\*) for grouping phenotypically similar lineages. We found that the uniform threshold *d*\*=2 gave visually well-defined clusters, and was consistent with the typical distance between well-measured lineages and their own coarse-grained gene knockouts (*d* ≃ 2 − 3). We used these definitions to search for knockout-like Tn lineages. For each of the strongly beneficial gene knockouts above, we computed Eq. (28) for ~20,000 candidate lineages that were measured in every environment and were located >100kb from the target gene. We deemed a lineage *ℓ_c_* to resemble the knockout of a particular gene if 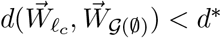.

After assembling a preliminary cluster of knockout-like Tn lineages for each target gene, we next checked that the clusters were likely to have been driven by secondary mutations, as opposed to the direct effects of Tn insertions in other functionally related genes. To be conservative, if an off-target gene was represented by multiple lineages in a given cluster, we removed those lineages from the cluster, since there is a chance that they could be driven by the direct effect of a Tn insertion in a gene that is functionally related to the target gene. This filter removed some lineages from the larger clusters like PUL66 (Fig. 4A), though we note that the number of multiply-represented genes in these clusters were consistent with a null expectation in which knockout-like lineages were randomly distributed across the genome. We additionally checked that no other beneficial gene knockout had a fitness profile within *d*\* of 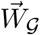 (after screening out the lineage in the other gene knockout that might have clustered with 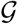).

The results of this pipeline are shown for two example genes in Fig. 4A,B; several others are shown in Fig. S13. Additional checks against noise-driven clustering and plotting details are discussed below.

#### Identifying novel clusters of secondary mutations

Loss-of-function mutations will likely comprise only a fraction of the adaptive landscape. Previous studies suggest that many of the adaptive lineages we detected in the main text may be caused by missense mutations and other noncoding variants (2), whose phenotypic effects—and therefore fitness profiles—may be qualitatively distinct from any gene knockout. To identify recurrent selection on such novel and strongly beneficial secondary mutations, we reversed the approach used in the previous section: we clustered individual lineages based on their own fitness profiles, and then looked for clusters of similar lineages that were different from every gene-level profile. We initially restricted this analysis to *Bc*, since this library was also measured in five additional *in vitro* environments.

To identify novel lineage clusters, we first curated a large list of Tn lineages that were well-measured and highly fit *in vivo*. We only considered lineages that had a measurable fitness in all environments (i.e. was present in at least one of the two time points comprising the environment), and had a strong positive fitness in at least one *in vivo* environment, 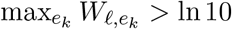. We applied bottom-up hierarchical clustering to the 2350 resulting lineages according to an average linkage criterion, using the distance metric Eq. (28) and a maximum merge distance of *ϵ* = 4. This yielded a total of 129 high-fitness clusters 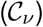, whose coarse-grained fitness profiles were defined by the unweighted average,

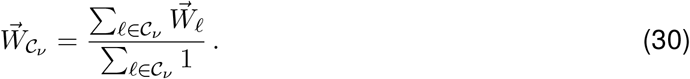

Since our initial high-fitness filtering omitted many adaptive lineages, we remapped the remaining 31,140 lineages with fitness measurements in all environments to their nearest high-fitness cluster. For each lineage *ℓ*, we found its distance-minimizing cluster 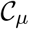:

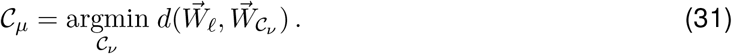

If this distance-minimizing cluster had *d* < *ϵ*, the lineage was added to the cluster. After adding 20,531 such lineages, the augmented clusters ranged in size from 1-2628 lineages, with a median of 29. Mean profiles were recalculated over the augmented clusters using Eq. (30).

We applied several stringent filters to discard clusters plausibly composed of lineages that acquired “knockout-like” secondary mutations (the subject of the previous section). First, we set aside 22 clusters with only one lineage: these could in principle be driven by a site-specific effect of its Tn insertion. We also excluded any cluster with fewer than 10 lineages that had 2 or more lineages in the same gene, since the probability of such a coincidence by chance is low (<1% per cluster). This removed only 4 clusters, consistent with our expectation from the main text that adaptation in *Bc* was not driven by gene knockouts. Using the same reasoning, we excluded 15 additional clusters that had more than 100 (or 1000) lineages and at least one gene represented more than 3 (or 4) times. This left a total of 94 candidate clusters.

We also directly compared the candidate clusters to gene knockout profiles. To focus on accurately measured gene profiles, we first compared the leave-one-out profiles (Eq. (29)) of each gene: if 2-means clustering isolated a single outlying profile, the corresponding lineage was removed and the gene profile was re-estimated. This removed much of the bias arising in genes harboring Tn lineages with strongly beneficial secondary mutations. As an additional control, we partitioned the remaining Tn lineages in each gene into two equal sized sets, and measured gene fitness profiles 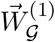 and 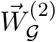 from the coarse-grained frequencies of the two subsets. We did not further consider the ≈ 35% of genes with unmeasured fitnesses in either subset. We then computed, for each cluster 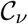, and gene 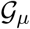, the distance 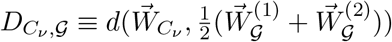 (excluding from 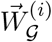 any lineage that was shared between 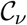 and 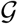). The use of the subset-averaged gene profile 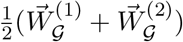 reduced the influence of any adaptive secondary mutations that had not been previously filtered out. Clusters with large values of 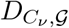 across all genes 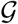 were deemed to be candidates for “novel” phenotypic clusters. Two examples, and their most similar genes (smallest 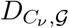), are shown in Fig. 4C,D in the main text; several more are shown in Fig. S14.

#### Controlling for ascertainment bias during clustering

Finally, for both novel and gene knockout-like clusters, we checked that our results did not arise from ascertainment biases arising from the clustering of thousands of noisy lineage profiles. We evenly split biological replicates in each environment into 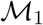 and 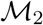, representing two independent sets of replicates across environments (analogous to the downsampling scheme employed above). For each lineage, we computed fitness profiles 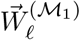 and 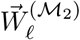, constituting independent measurements of the fitness profile that should be similar to one another, as well as to other 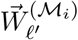 in the cluster. We calculated the unweighted average of the two replicate profiles, 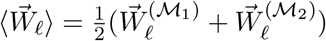, for each lineage. The interquartile range of 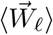 among co-clustered lineages is represented by the green shaded bands in Fig. 4 and Figs. S13 and S14. (If a cluster had fewer than 10 lineages, the full range is plotted instead.) Similarly, the black shaded regions in Fig. 4 and Figs. S13 and S14 represent the spread between 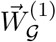 and 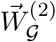, described in the previous subsection.

### 7 Evidence for additional *de novo* mutations

The dynamics we observed in the main text were dominated by lineages that exhibited consistent fitness benefits in independent mice, implying that the causative mutations were already present in the library before gavage (4, 19). However, it is plausible that additional adaptive mutations continued to accumulate within these populations during the subsequent two weeks of the experiment (1, 4). Such a mutation would initially arise within a single Tn lineage within single a mouse. If the mutation survived genetic drift, it would sweep through its Tn lineage and continue to spread through the population, leading to a lineage trajectory that diverged from the other biological replicates.

To examine the evidence for such mutations, we ranked the day 16 frequencies of each lineage ℓacross all *k* = 5 mice in the HF/HS diet, *f*_*ℓ*,1_(16) ≥ *f*_*ℓ*,2_(16) ≥ … ≥ *f*_*ℓ,k*_(16). We defined a divergence metric Δ_*ℓ*_ as the ratio between the largest and second largest frequencies,

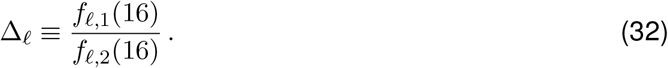

Large values of Δ_*ℓ*_ would be consistent with a beneficial mutation occurring in mouse 1. However, these divergent lineage trajectories could also arise through biological or technical noise (e.g. if a lineage was measured at small read counts across mice). To mitigate the impact of these effects, we restricted our attention to lineages that reached >0.1%frequency in at least one mouse (which required that they were sampled in at least 10 reads).

Among these lineages, we observed a clear enrichment in large divergence values (Δ_*ℓ*_ > 10) in *Bo*, *Bt-VPI*, and *Bt-7330*, but comparatively few divergent lineages in *Bc* (Fig. S15). In the first three species, these large divergence lineages accounted for 20%-75% of all lineages that reached >0.1% frequency. The trajectories of five randomly chosen examples from each species are shown in Fig. S16. These randomly chosen examples are highly suggestive of de novo adaptive mutations, in which the large divergence between the largest and next-largest lineage is maintained across multiple sequenced timepoints. Furthermore, while large divergence values could also be produced by the stochastic establishment of pre-existing mutations, we found that most of the examples in Fig. S16 were not consistent with this behavior, since the remaining lineages tended to maintain or decline in frequency over time. Interestingly, many of the divergent trajectories in Fig. S16 grew most rapidly over days 4-10, and grew more slowly (or even declined) over days 10-16. This could arise from further shifts in selection pressures during days 10-16, or simply from increases in the mean fitness 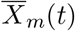. Together, these examples suggest that there is at least some evidence for adaptive *de novo* mutations during the first 16 days of the experiment. Future experiments over longer time periods and with higher temporal resolution will be necessary to fully resolve the interplay between *de novo* mutations and standing genetic variation.

### 8 Estimating the strength of genetic drift *in vivo*

Our results in the main text suggest that positive selection is a pervasive force during *in vivo* colonization (Figs. 1–3). The opposing force of genetic drift could also play an important role *in vivo*, reflecting both the enhanced spatial structure within the gut, or transient population bottlenecks during engraftment. Here, we describe a method to measure the strength of these genetic bottlenecks when natural selection is widespread. We apply this algorithm to simulations and experimental data.

In the absence of natural selection 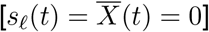, genetic drift leaves a well-known signature in the variance of a lineage’s frequency,

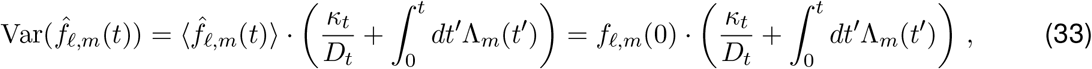

which follows from the model in Eq. (9). This simple behavior underlies several common methods for inferring the strength of genetic drift from frequency trajectory data (45–49). In the presence of natural selection, this behavior takes on a more complicated form,

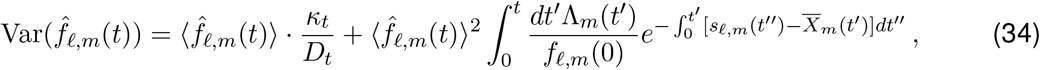

which depends on the fitness of the focal lineage (*s_ℓ_*(*t*)) as well as the mean fitness of the population 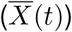. Estimating the strength genetic drift in this more general scenario is more challenging.

Previous work has shown that the strength of genetic drift can be inferred from a collection of neutral marker lineages (*s*ℓ**(*t*) = 0) with a sufficiently dense time series (19, 20). However, this approach suffers from two key limitations that make it difficult to apply in our present case. First, if a substantial fraction of the marker lineages have non-neutral fitnesses, this method will be strongly biased by the variation in fitness among the marker lineages, which will tend to overestimate the strength of genetic drift. Second, this approach also requires multiple sequencing replicates at each timepoint to distinguish the contributions of genetic drift from technical noise (19).

Here we sought to exploit a different feature of Eq. (34) to infer the strength of genetic drift when natural selection is sufficiently widespread. Our approach is based on the observation that contribution from technical noise depends on the present-day frequency of the lineage, while the contribution from genetic drift depends on the historical trajectory of the lineage as well. For a given present-day frequency, a higher fitness lineage must have been present at a lower frequency in the initial timepoint, and will therefore have experienced stronger genetic drift on the way to its present-day frequency. This suggests that the strength of genetic drift will show up as a systematic correlation between 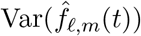 and 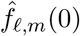.

To make this intuition more precise, we focused on a simple version of the model in Section 4, in which the relative fitnesses and strength of genetic drift are approximately constant over the relevant time interval. In particular, for every mouse *m* with host environment *e*, we let *χ_ℓ,e_*(*t*) ≈ *χ_ℓ,e_* and define Λ_*m*_(*t*) ≃ 1/*N_e_τ_e_*, where *N_e_* is the effective population size and *τ_e_* is the effective generation time in the gut. In this limit, Eq. (34) reduces to

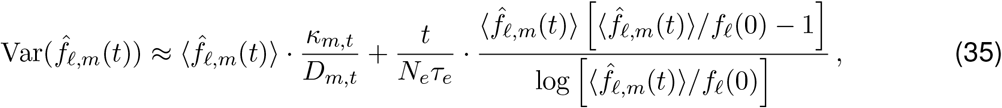

where

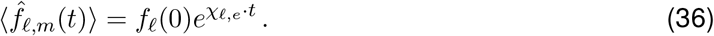

Since the average frequency is independent of *m*, the variance can be estimated from the divergence in the lineage’s frequency across mice 1 and 2:

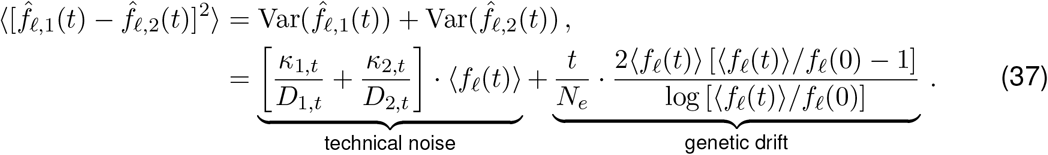

This suggests that the contributions from genetic drift and technical noise can be distinguished by their different scaling as a function of 〈*f_ℓ_*(*t*)〉 and *f*_*ℓ*_(0). In particular, if we define the predictor variables

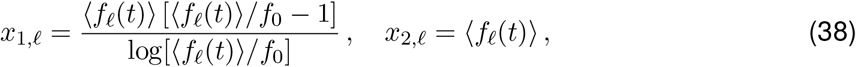

then the magnitude of genetic drift (*t*/*N_e_*) can be inferred from a linear regression of 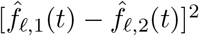 on *x*_1,*ℓ*_, and *x*_2,*ℓ*_.

However, a straightforward implementation of this regression approach is challenging, since the predictor variables must also be self-consistently estimated from the data. The additional uncertainty in these estimates leads to substantial and heteroscadastic noise in the underlying regression model that will bias the estimated regression coefficients (50). To mitigate these issues, we instead performed regression on pre-averaged groups of *k* ≫ 1 lineages, which were chosen to have similar initial frequencies and relative fitnesses. The inputs to the regression model are the average values of the predictor and response variables within each group *L*:

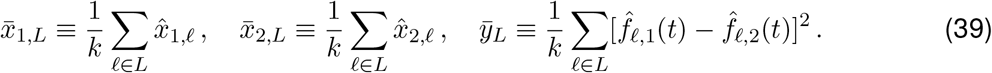

We estimated the individual contributions 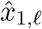 and 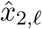 using the plug-in estimators,

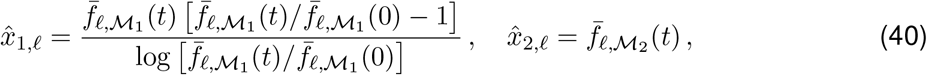

where 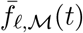 denotes the cohort-averaged frequency defined by Eq. (16b), and 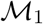 and 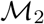 are non-overlapping cohorts that do not contain mice 1 or 2. We also assigned each group of a lineages a corresponding weight

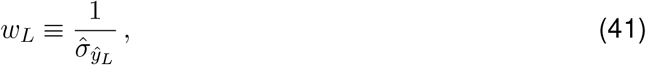

where 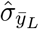 is the standard deviation of *ȳ_L_* estimated by bootstrap resampling within *L*. A weighted regression of *ȳ_L_* on 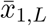 and 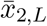 yields a corresponding estimate for *t*/*N_e_τ_e_*.

As a proof of principle, we applied this algorithm to measure genetic drift in the *Bc* population, which benefited from strong fitness variation and weaker sequencing noise compared to the other *Bacteroides* species. We first tested the algorithm on a simulated set of biological replicates designed to mimic the conditions of our experiments (as described above to validate rank-order curves): we generated an empirical distribution of lineage fitnesses during days 0-4 as described in SI Section 3. These inferred fitnesses, along with the input frequencies, were used to initialize 9 identical populations. We simulated these populations for 40 generations (~4 days) under a Wright-Fisher model with a fixed generation time of *τ_e_* = 10/day and population size *N_e_* varying from 10^5^ – 10^9^ days. Finally, to generate simulated sequencing libraries, we modeled the sampling and sequencing of each population as a sequence of two Poisson sampling steps, each equal to the empirical sequencing depth at day 4 in the corresponding mouse.

We then applied our drift inference algorithm to these simulated populations. We first used the input frequencies and simulated sequencing libraries to partition the lineages into groups of at least 50 lineages with similar initial frequencies and average fold changes across mice. (To mimic the conditions of the experiment, these average fold changes were estimated from the simulated sequencing libraries, rather than the true fitnesses in the simulation). Given these lineage groupings, we partitioned the biological replicates into three non-overlapping cohorts and used them to estimate 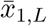 (4 replicates), 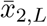 (3 replicates), and *ȳ_L_* (2 replicates) using Eqs. (39) and (40). The effective population sizes (*N_e_τ_e_*) estimated from the linear regression are shown in Fig. S17. We observed that over different permutations of mice in the three cohorts, our algorithm could reliably distinguish population sizes over several orders of magnitude, unless genetic drift contributed <1% of the variance attributable to sequencing noise.

Having validated our algorithm on simulated data, we applied it to the 9 mice that were subject to the HF/HS diet over days 0-4 (Fig. S17). The effective population size estimate in this time interval was around 5 · 10^6^ days, which coincided with the regime in which there was good agreement between the inferred values and the ground truth in our simulations. Our estimates suggest that genetic drift accounted for only ~10% of the observed variance in lineage frequencies across replicates, highlighting the potential of our approach to distinguish the effects of biological noise even when technical noise is relatively strong. Interestingly, this estimate is substantially lower than the census population sizes estimated for *Bacteroides* species in the murine gut (25).

However, these initial estimates should also be treated with a degree of caution, since our simple regression model made a number of simplifying assumptions that may not hold in practice. Chief among these was the assumption that Λ(*t*) and *χ_ℓ_*(*t*) are approximately constant over the relevant time interval, though residual noise or co-linearity in the predictor variables could also pose problems for the linear regression step. The reasonable performance of our algorithm on simulated data suggests that these issues have a limited impact in the parameter ranges we examined here. Further refinements to this algorithm – and extensions to other sources of biological noise – would be an interesting topic for future work.

**Figure S1:**
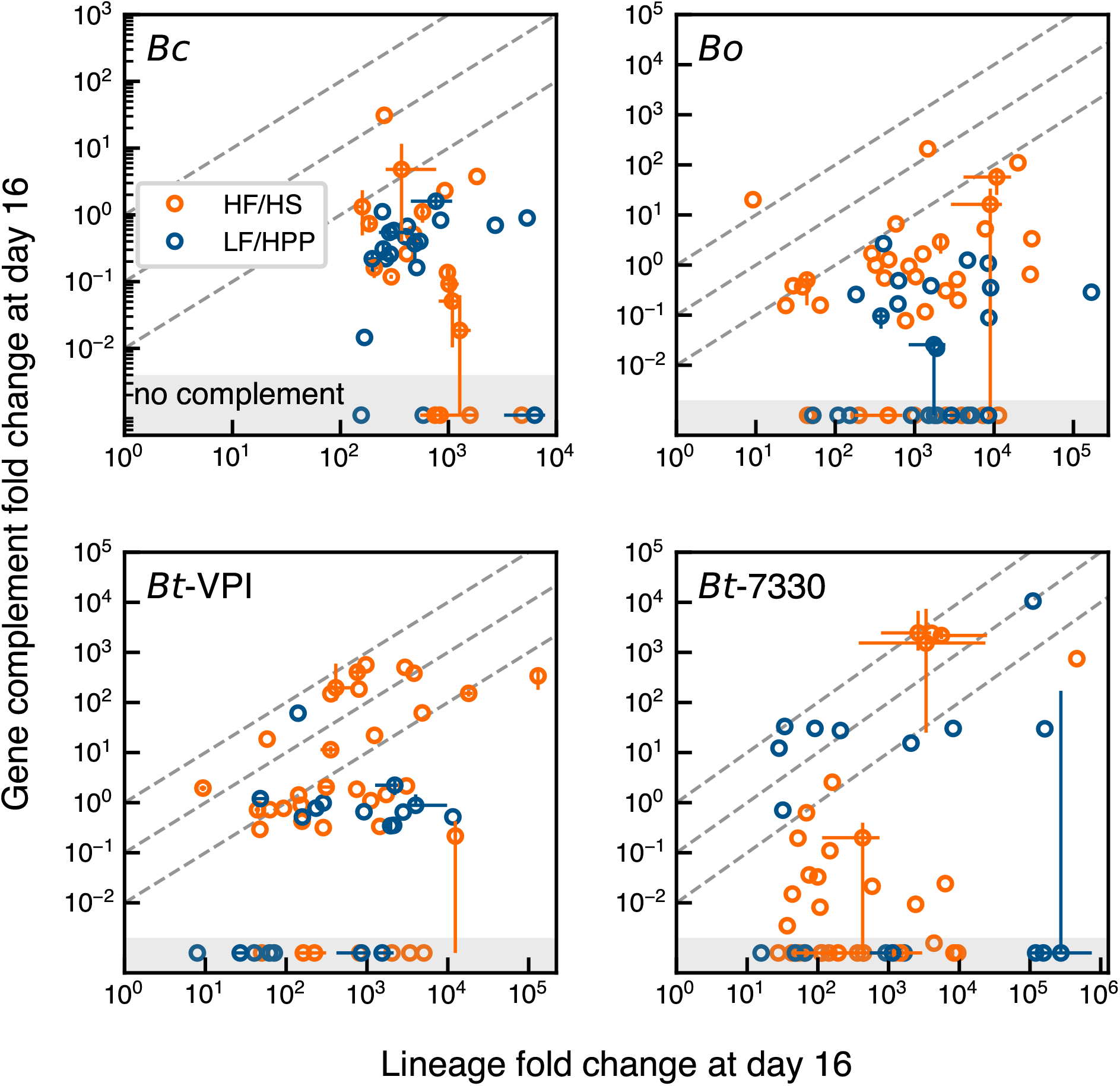
Most lineages that reached intermediate frequency by day 16 were not driven by beneficial Tn insertions. For each mouse in the HF/HS (red) or LF/HPP (blue) diets, the fold change of the top 10 largest lineages at day 16 is plotted against the of other Tn lineages in the same gene (the gene complement, SI Section 4). Lineages among the top 10 in multiple mice in the same diet are represented by their median (circles) and minimum and maximum values (lines) across those mice (lines). Lineages in the grey region either have Tn insertions in intergenic regions or gene complements measured at zero frequency at day 16.

**Figure S2:**
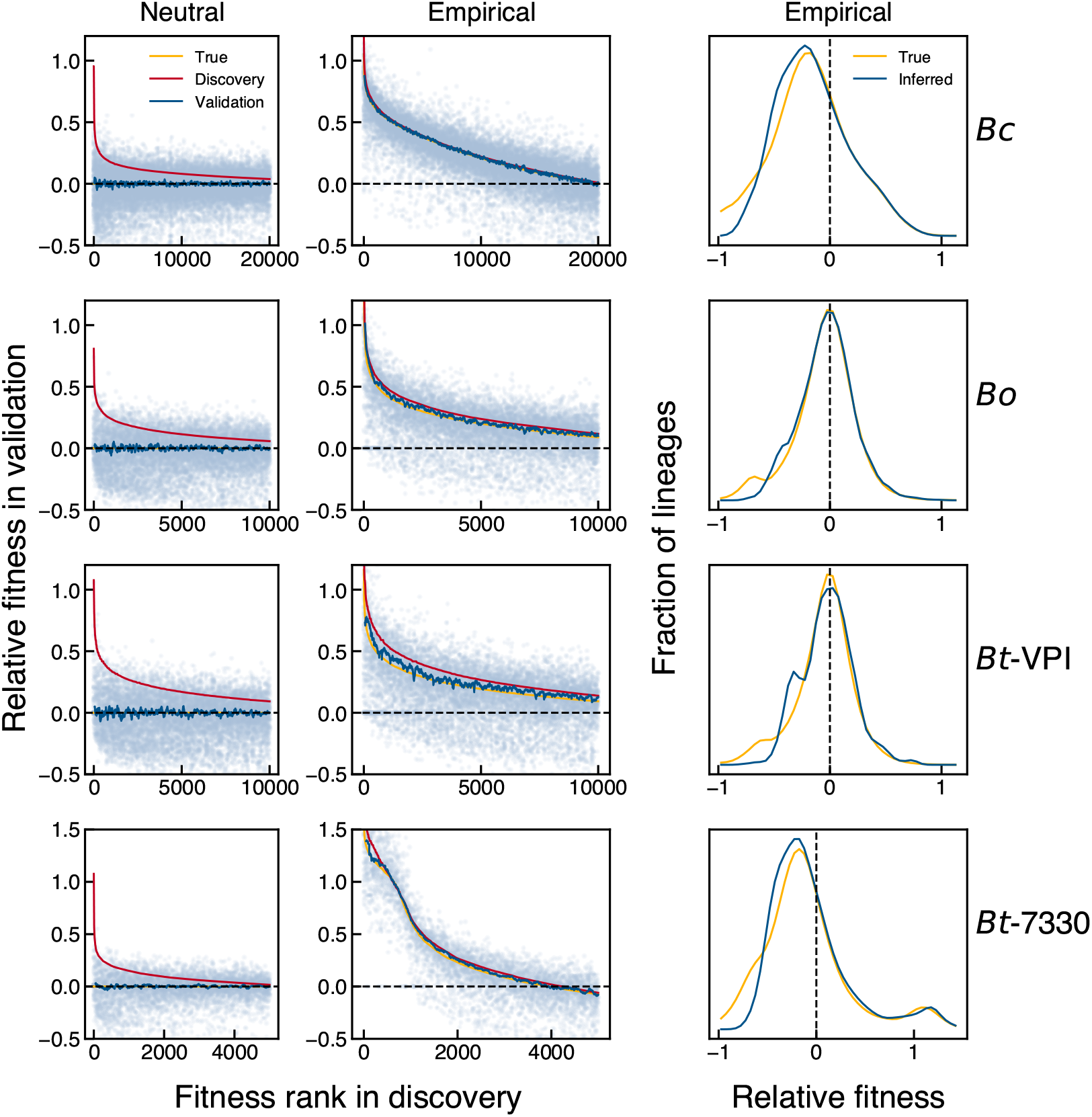
Cross-validation consistently estimates relative fitnesses of lineages in simulated populations. For each *Bacteroides* library (row), 9 populations were initialized with the distribution of lineage frequencies estimated from the pooled day 0 input libraries. In one set of simulations (left column), every lineage’s fitness was set to 0 to mimic a neutral scenario. In another set (center column), each lineage’s fitness was set to its average across 9 HF-fed mice during days 0-4, according to Eq. (16). Lineage dynamics were simulated as described in SI Section 4. Lineages were then filtered, ranked, and plotted as described in SI Section 4. The right plot compares the true distribution of lineage relative fitnesses to that estimated by inverting the coarse-grained rank-order curve in the center plot (SI Section 4).

**Figure S3:**
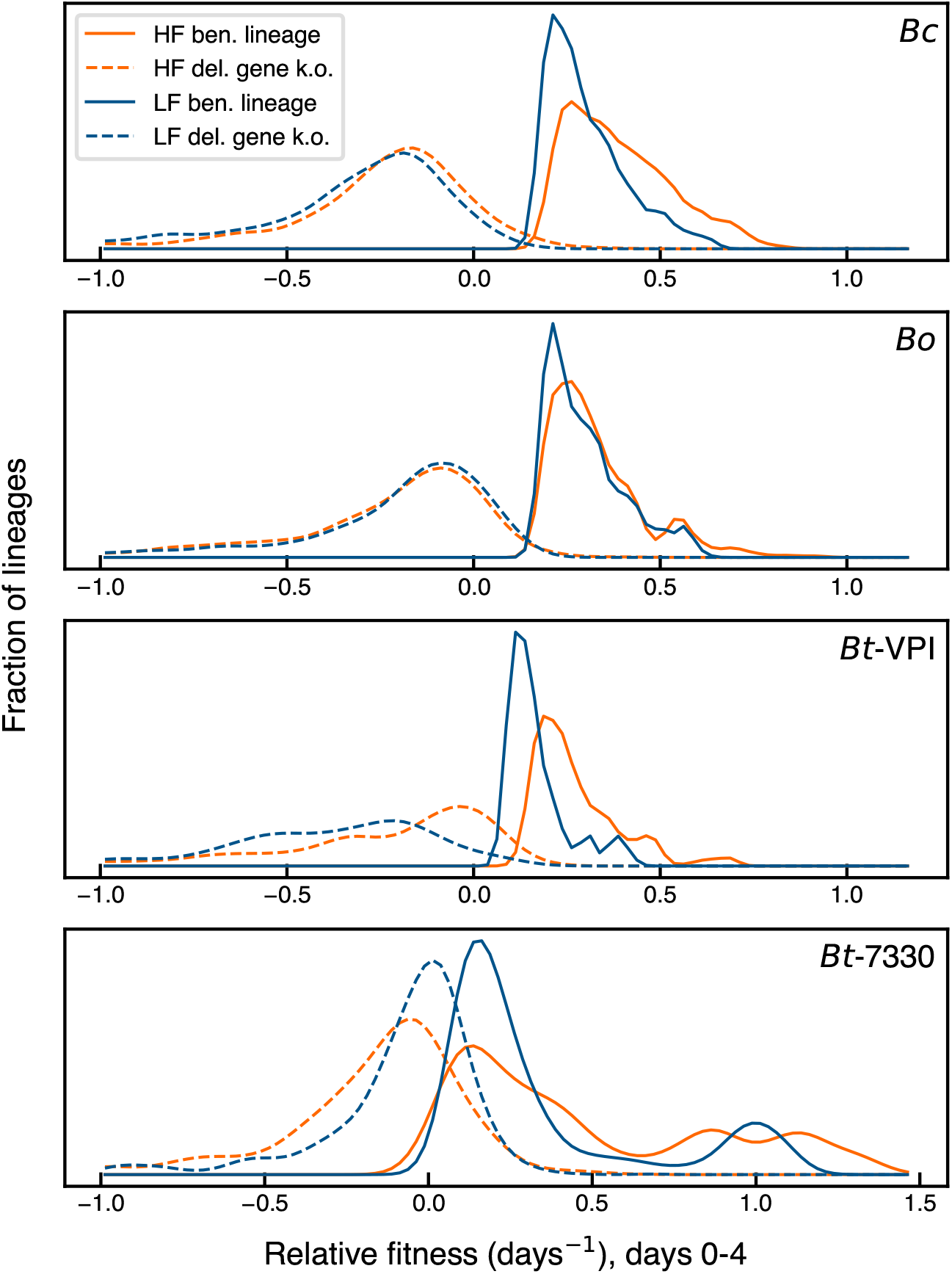
Relative fitnesses of adaptive lineages compared to “fitness determinant” gene knockouts identified in previous work. For each *Bacteroides* library, the relative fitnesses during days 0-4 of deleterious gene knockouts, previously annotated by Wu *et al*. (28), were estimated in a cohort of mice. Frequencies of gene-knockouts were estimated by summing all reads from Tn insertions falling in the gene, as defined in SI Section 4. Finally, Gaussian kernel density estimation was applied to plot a smoothed distribution. This distribution was compared to cross-validated relative fitnesses, measured in the same cohort of mice and over the same interval, of the fittest 10,000 (*Bc*), 5,000 (*Bo* and *Bt-VPI*), or 3,000 (*Bt-7330*) lineages, which were estimated with a separate discovery cohort of mice. The smoothed distribution of adaptive lineages was computed as described in SI Section 4.

**Figure S4:**
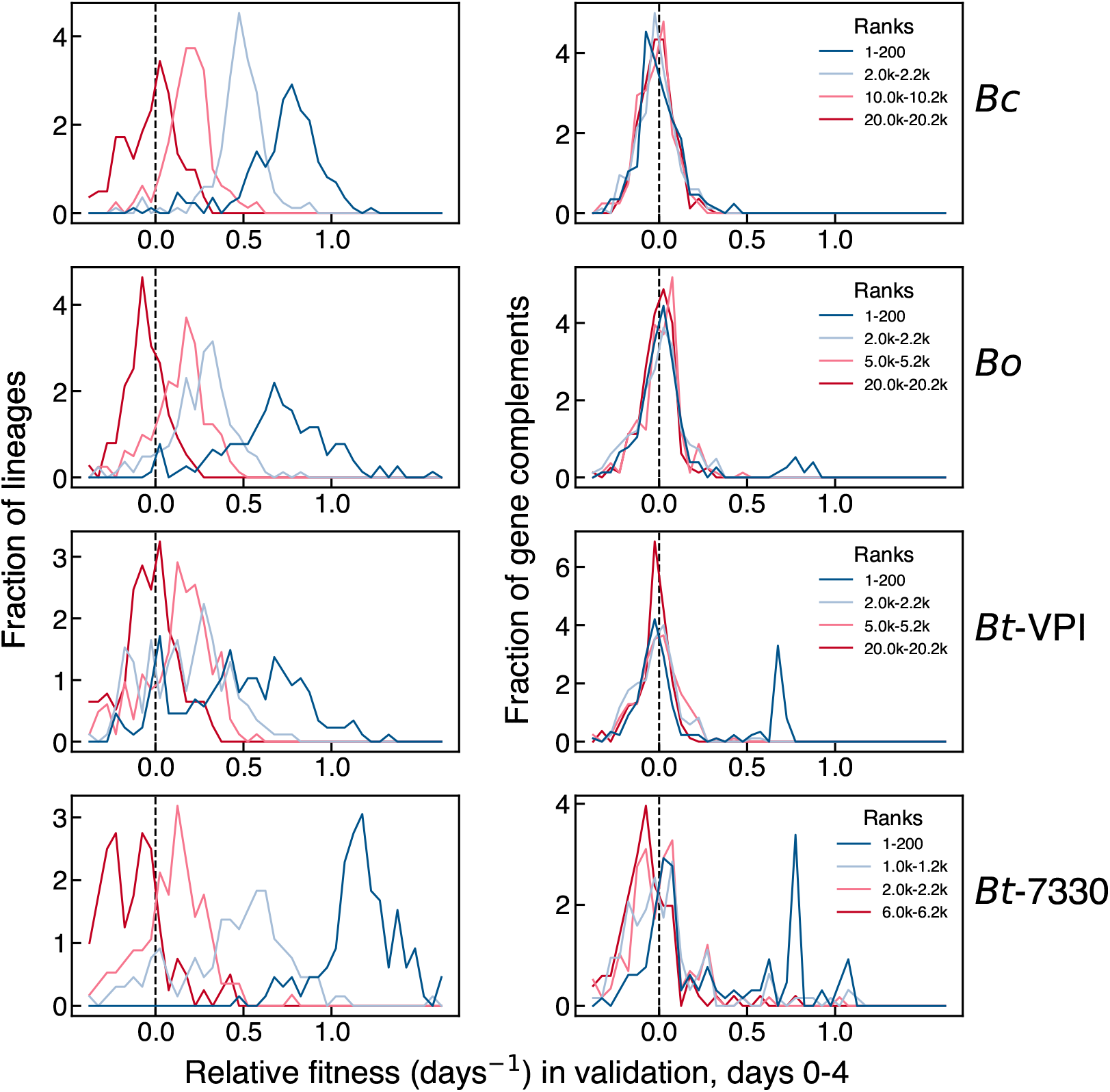
Conditional distributions of relative fitnesses at different ranks. In each *Bacteroides* library (**row**), 4 sets of lineages were collected, spanning different ranks of relative fitness during days 0-4 in the HF/HS discovery cohort. For each set, the distributions of relative fitnesses of the lineages (**left**) and their gene complements (**right**, SI Section 4) in the HF/HS validation cohort during days 0-4 are plotted. The same cohorts of HF/HS discovery and validation mice were used as in Fig. 2 and Fig. S5. In each of the 4 *Bacteroides* libraries, the mode of adaptive lineages’ relative fitness remains positive over 1000s of ranks, suggesting positive fitness variation across 1000s of lineages. Conversely, the mode of their respective gene complements’ fitnesses are centered at 0 relative fitness, independent of rank, suggesting that most lineages’ fitness benefits do not derive from the gene-knockout effects of their original Tn insertions.

**Figure S5:**
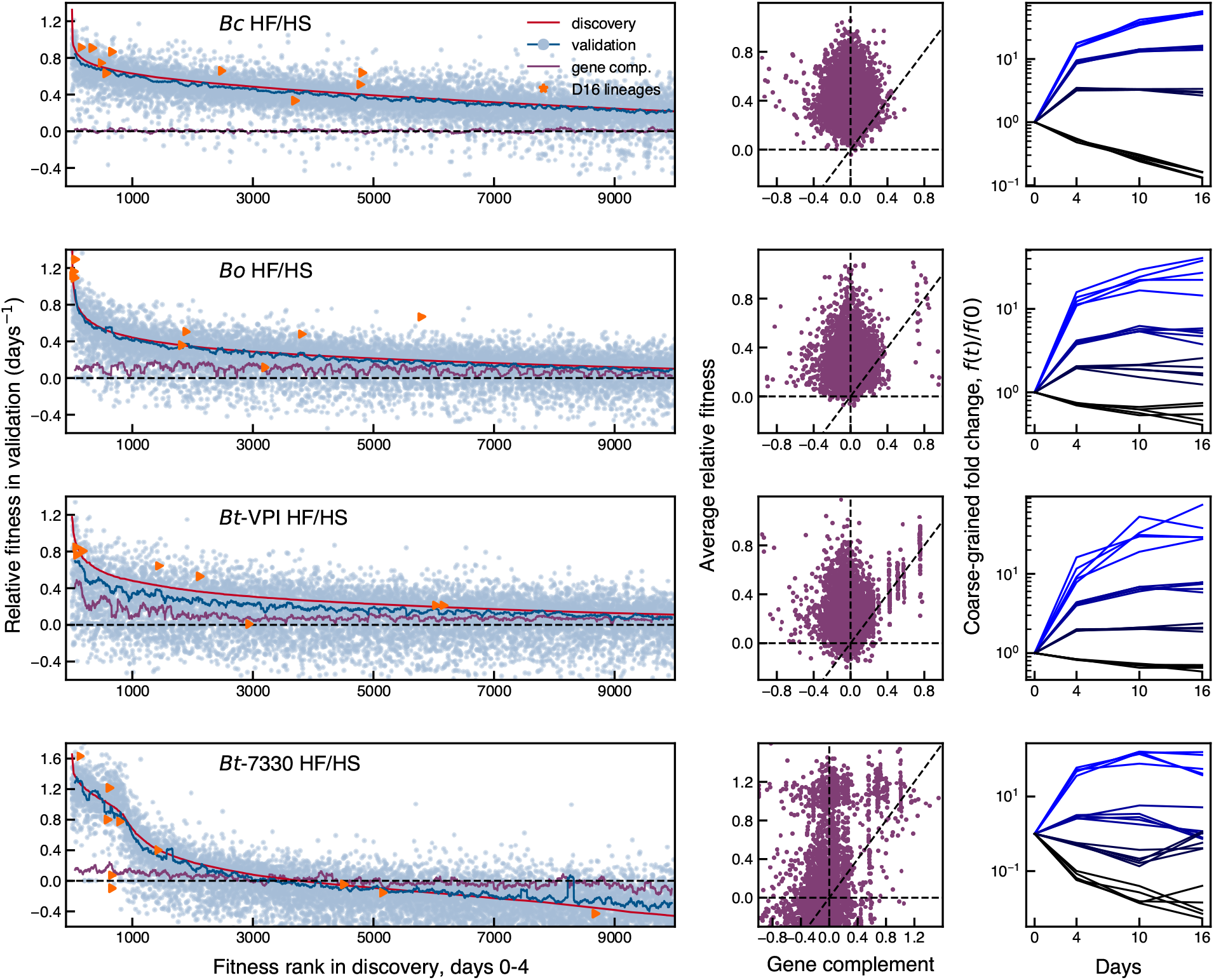
Analogous version of Fig. 2C-H for all four species. The same sets of discovery and validation mice are used as in Fig. 2. To emphasize the difference in the numbers and fitnesses of adaptive lineages, the first 10,000 ranks are shown for all 4 *Bacteroides* species. In the rightmost plots, colored lines show the total frequency over time of ranks 1-1,000, 1,000-3,000, and 3,000-10,000 (light blue to dark blue), as well as the remaining lineages (black) in different mice.

**Figure S6:**
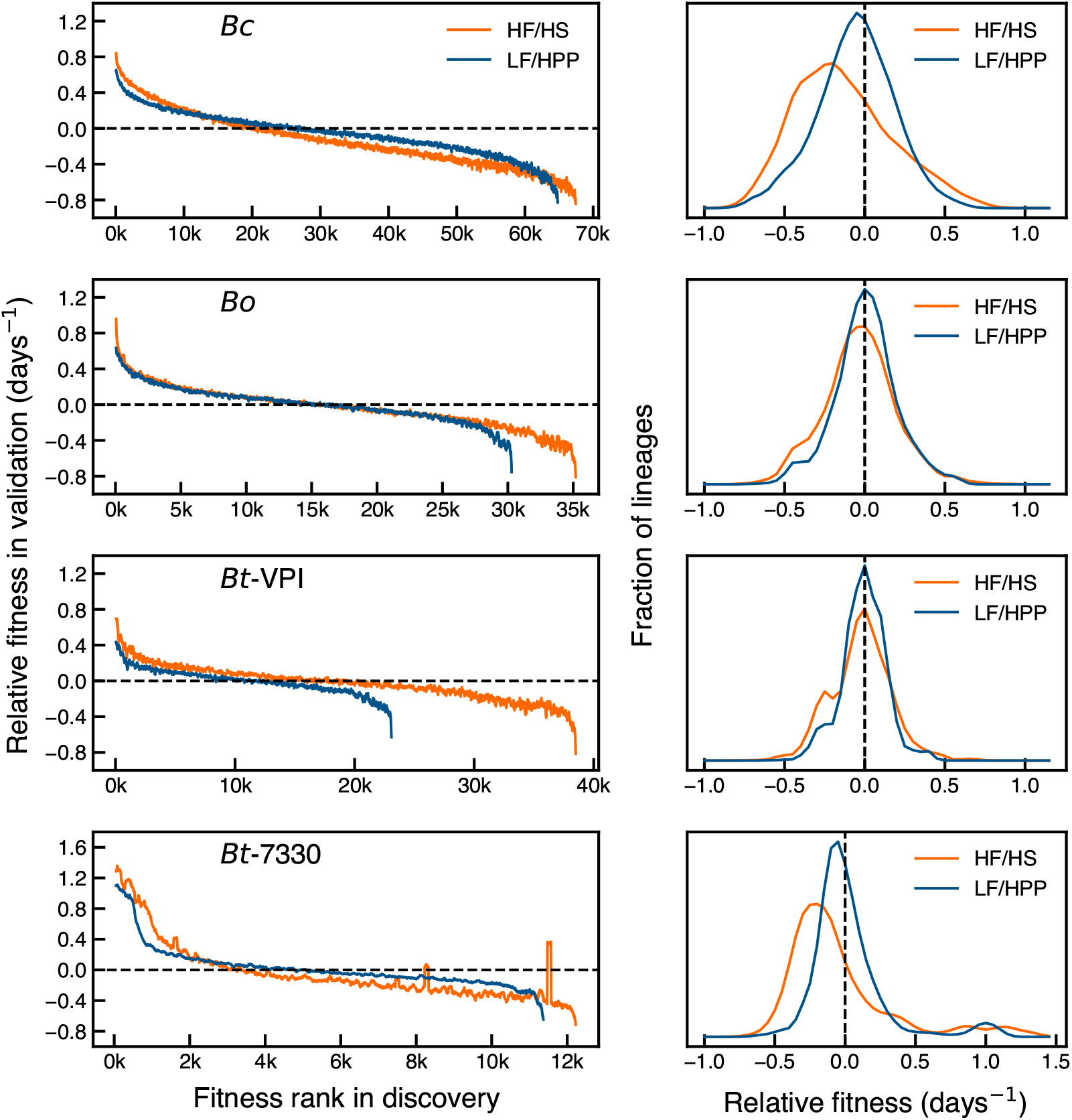
Full rank-order curves and distribution of relative fitnesses in the HF/HS and LF/HPP diets. For the HF/HS diet, 5 mice were used for discovery and 4 mice were used for validation, while in the LF/HPP diet, 4 mice were used for discovery and 3 were used for validation. Note that different sets (and numbers) of lineages pass the filtering steps in each diet (SI Section 4).

**Figure S7:**
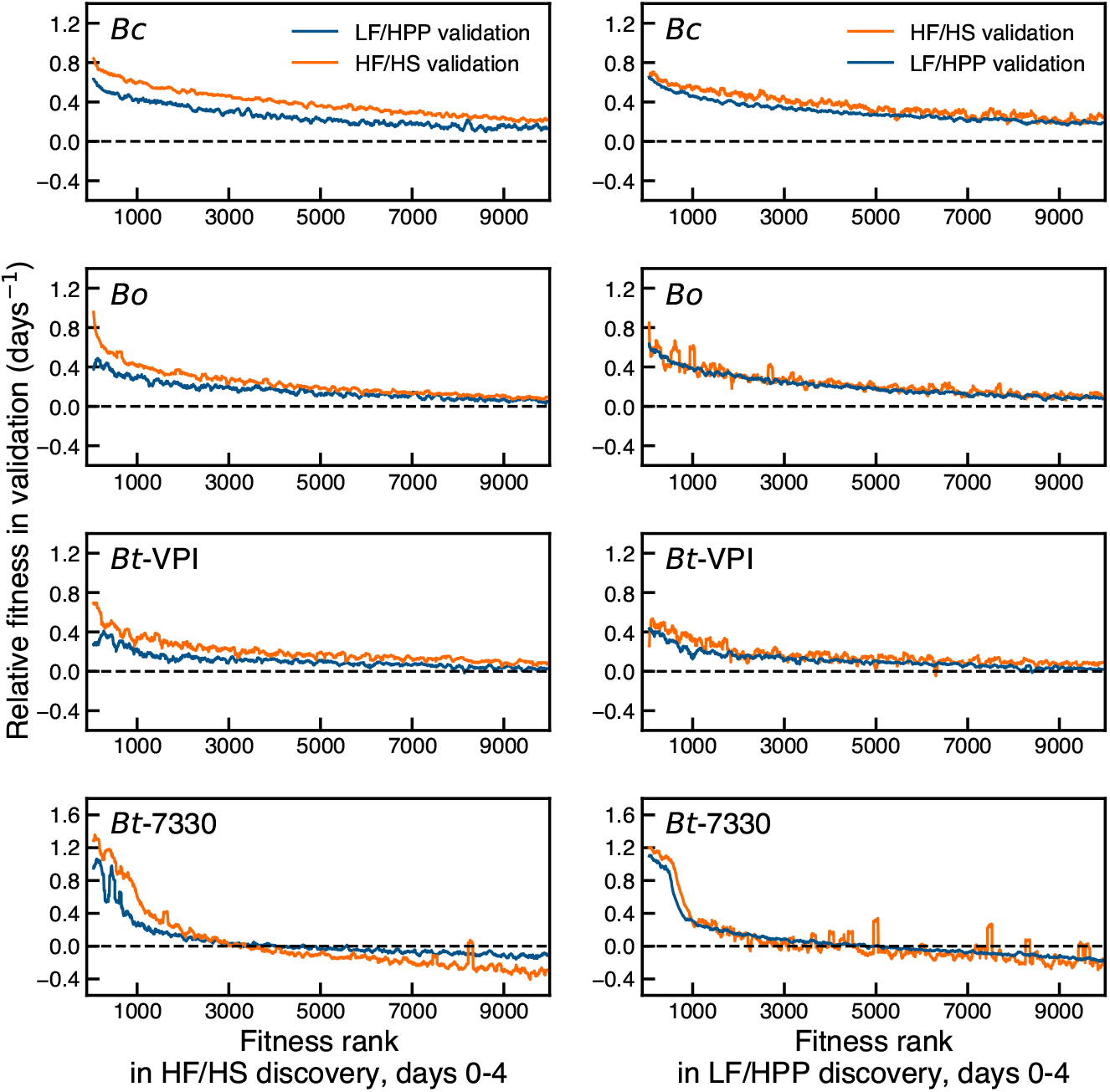
Lineage fitnesses are strongly correlated across diets during days 0-4. Relative fitnesses during days 0-4 in the HF/HS and LF/HPP diets plotted against rank order of relative fitnesses during days 0-4 in a discovery cohort with the same (blue) or different (red) diet. Each curve represents a running average of 100 lineages.

**Figure S8:**
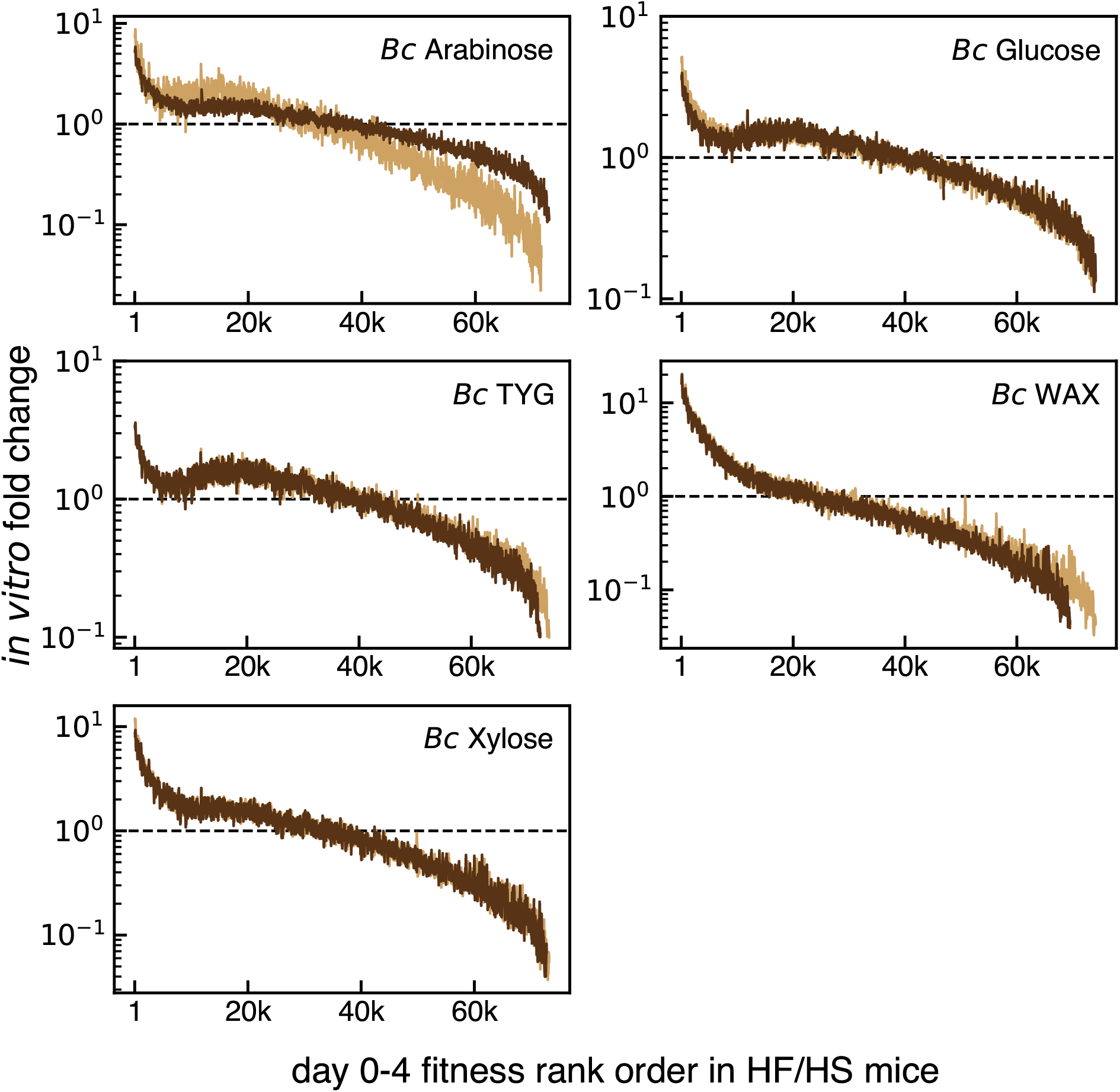
*In vivo* relative fitnesses correlate with *in vitro* relative fitnesses for a subset of media in *Bc*. Fold changes of *Bc* lineages in overnight cultures of different media, ranked by their relative fitnesses during days 0-4 in HF/HS mice. The curves represent running averages of 100 ranked lineages (SI Section 4). For each medium, two independent cultures (gold, brown) demonstrate the reproducibility of the relative fitnesses of coarse-grained lineages.

**Figure S9:**
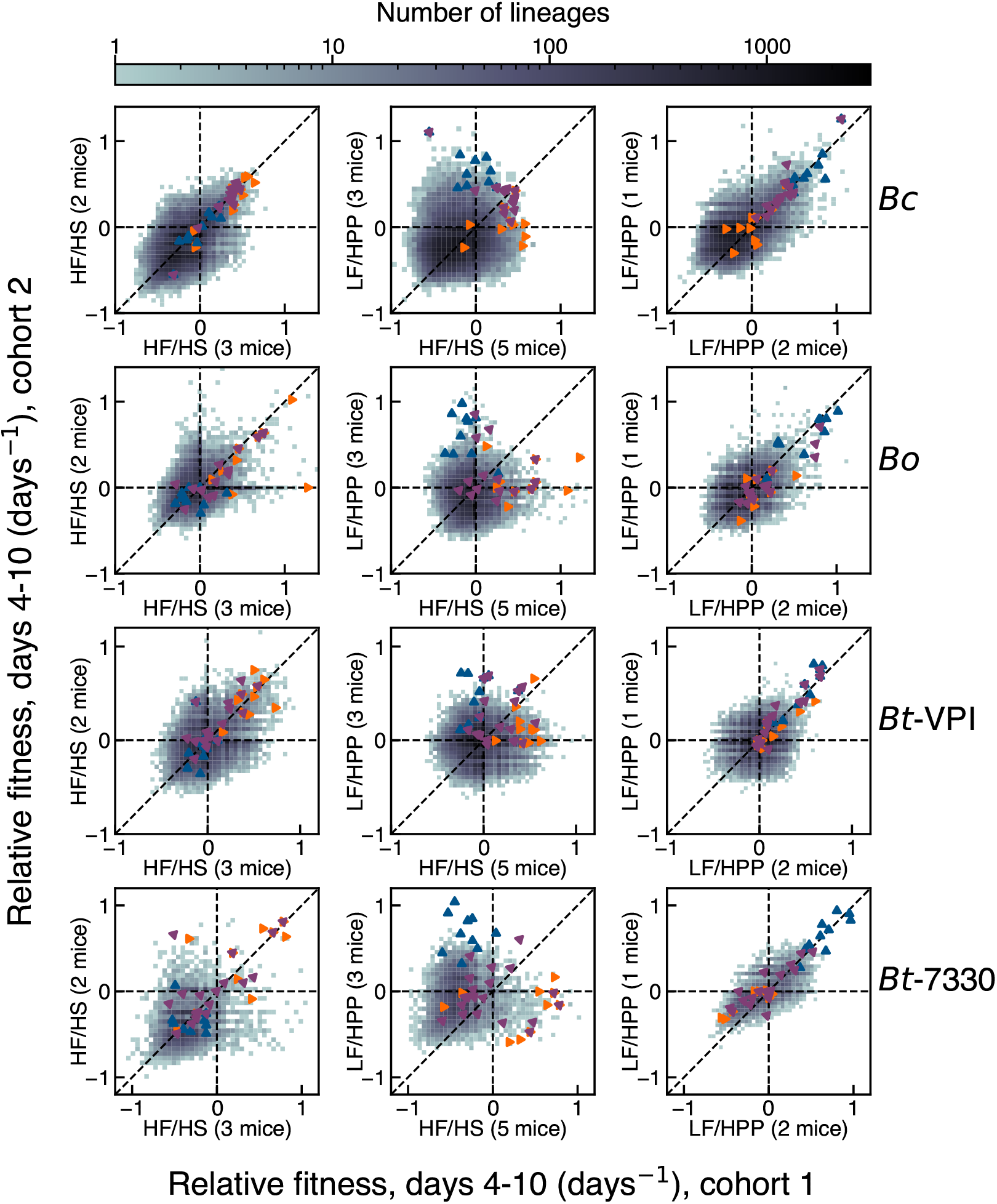
Lineage fitnesses during days 4-10 are correlated within but not between diets. For each panel, average relative fitnesses during days 4-10 were calculated in nonoverlapping sets of HF/HS and LF/HPP mice (SI Section 4). Rows correspond to different *Bacteroides* species and columns to the same sets of mice. Triangles indicate the 10 largest lineages at day 16 in the HF/HS (orange), LF/HPP (blue), or alternating (purple) diets.

**Figure S10:**
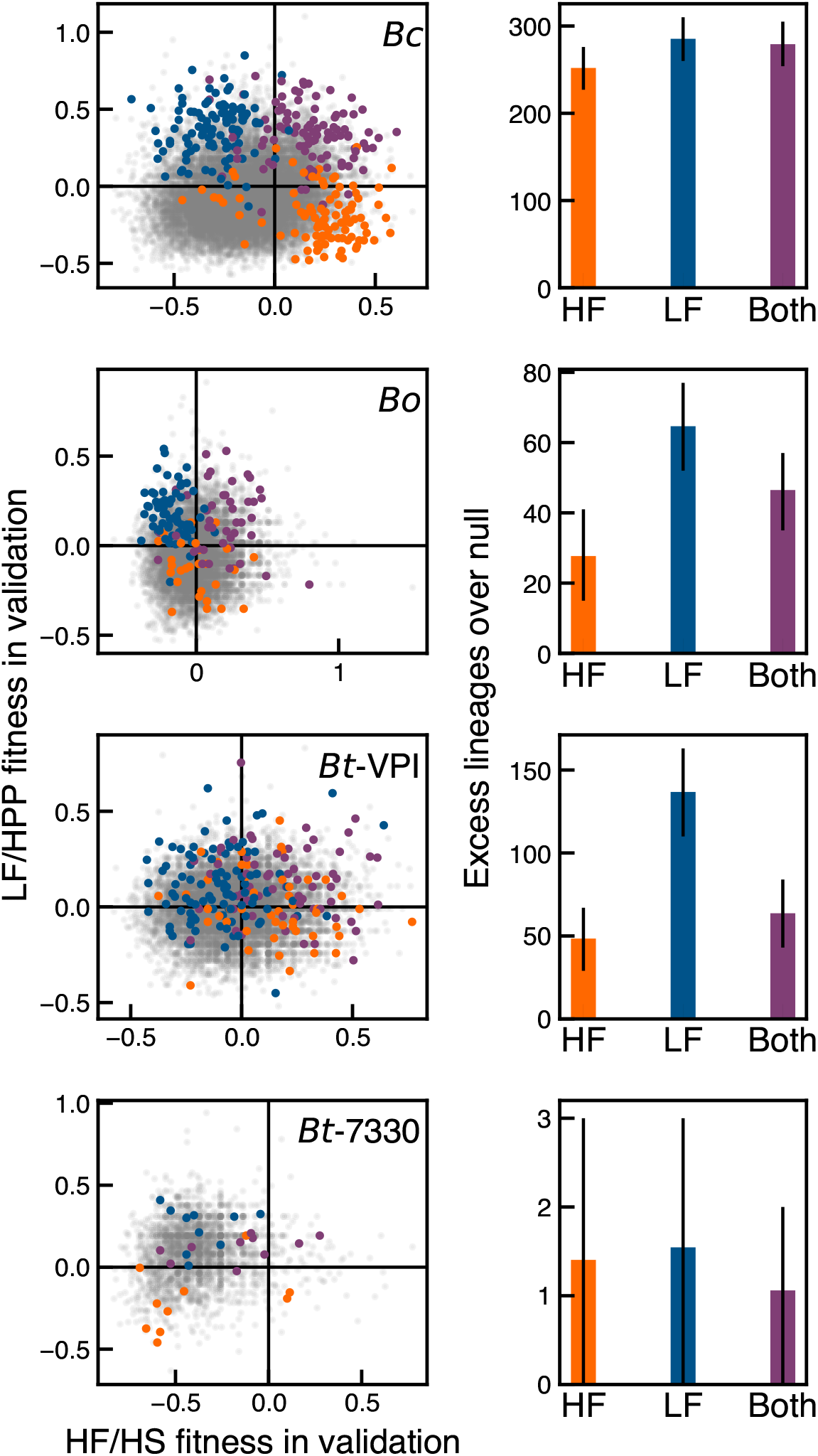
Hundreds of lineages exhibit relative fitnesses that depend on diet at later times. For each *Bacteroides* species, the left panel shows the joint distribution of all relative fitnesses between days 4-10 (grey) in the a cohort of 2 HF/HS and 1 LF/HPP mice. The highlighted lineages have the largest value of the tradeoff statistic |*T_ℓ_*| in a separate discovery cohort of 3 HF/HS and 2 LF/HPP mice (SI Section 5). The right panel shows the excess number of lineages, over a null expectation (SI Section 5), in each quadrant in the discovery cohort that fell in the same quadrant in the validation cohort (dark shading). Error bars indicate 95% confidence intervals in the excess lineages, from repeated draws of the null distribution.

**Figure S11:**
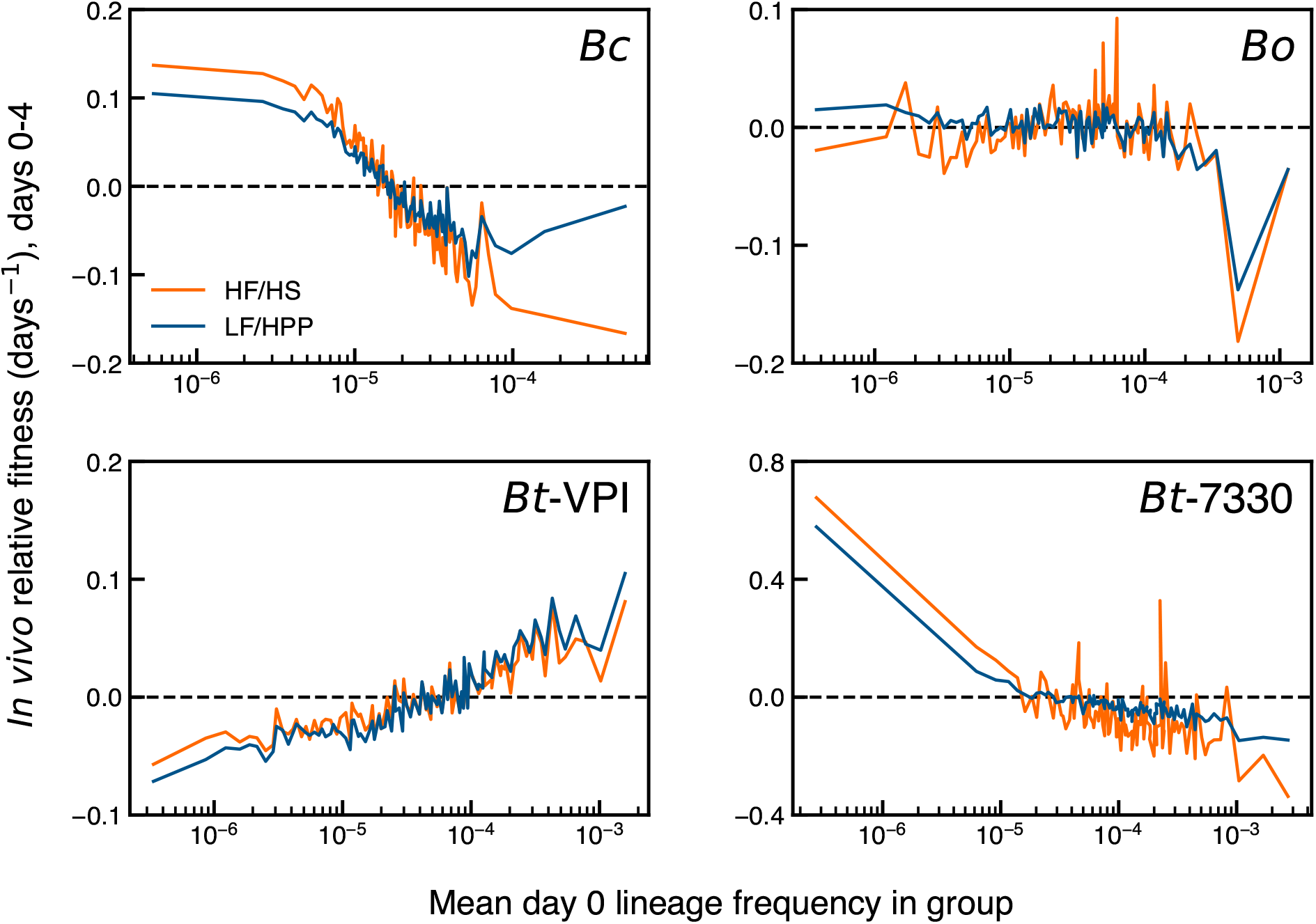
Lineage fitnesses are anti-correlated with their frequency in the input library in *Bc* and *Bt-7330*. For each species, day 0 lineage frequencies were estimated and ranked in a discovery pool of input libraries. Lineages were then coarse-grained into 100 super-lineages of roughly equal day 0 frequency ~1% using the procedure described in SI Section 4. The relative fitnesses of these super-lineages were estimated during days 0-4 *in vivo* using a separate validation pool of input libraries to re-measure initial frequencies. Relative fitnesses in HF/HS (orange) and LF/HPP (blue) cohorts of mice are plotted against the average lineage frequency within each super-lineage, measured in the validation input pool. *Bc* and *Bt-7330* exhibit strong anti-correlations between *in vivo* fitness and initial frequency, whereas *Bt-VPI* shows the opposite trend.

**Figure S12:**
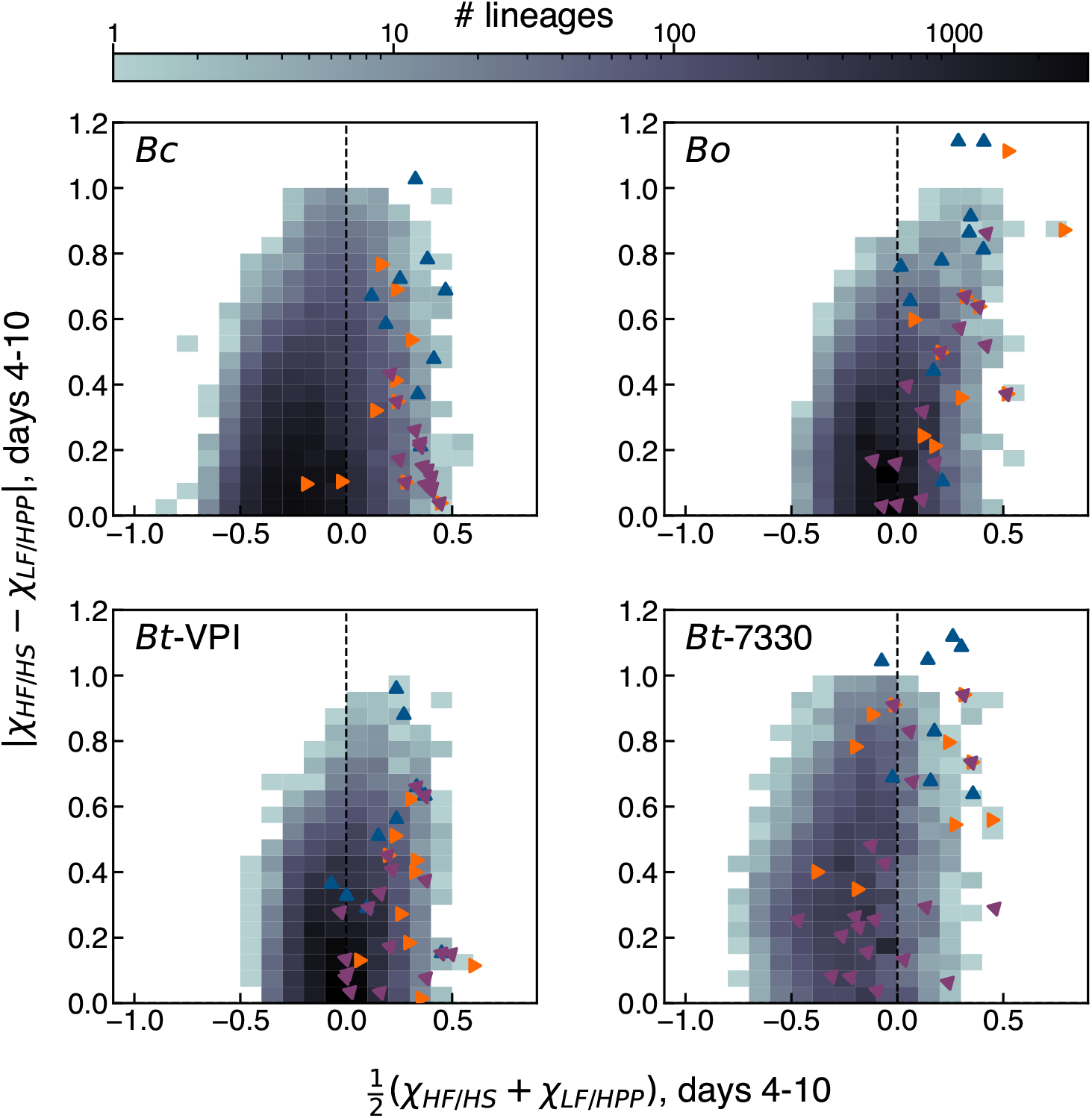
Analogous version of of Fig. 3C computed for diet-averaged fitnesses. For each of the lineages in Fig. 3C, we computed a diet-averaged fitness 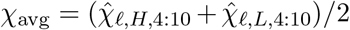 and the corresponding off-diagonal component 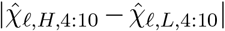 between days 4-10. An analogous calculation was carried for the other three species. This projection shows that the largest lineages in the alternating diets (purple triangles) had higher diet-averaged fitness and smaller tradeoffs in *Bc*, despite their smaller representation in the underlying distribution.

**Figure S13:**
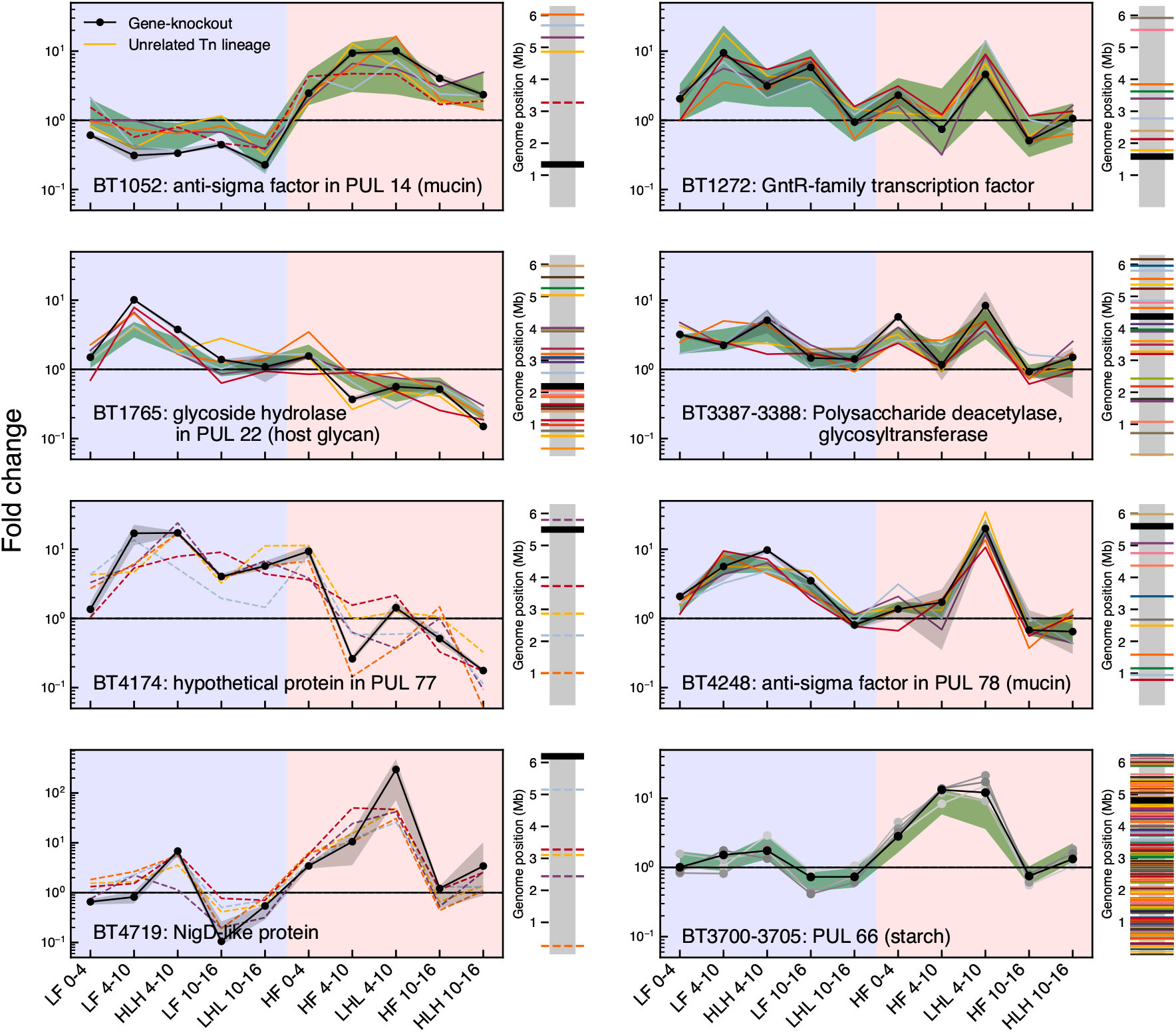
Analogous version of Fig. 4A,B. Strongly beneficial *Bt-VPI* gene knockouts (black curve) and unrelated Tn lineages with similar fitness tradeoffs (colored curves and green band; SI Section 6). For gene knockouts with fewer than 5 unrelated Tn lineage within a fitness profile distance *d* < *d*\* = 2 (Eq. (28)), the 5 lineages with the smallest distances are plotted; dashed lines denote lineages with *d* > 2. At bottom right, 235 Tn lineages cluster with gene knockouts of BT3700-BT3705 (excluding BT3704); the fitness profiles of these genes (shades of grey and black) are shown to highlight their consistent behavior.

**Figure S14:**
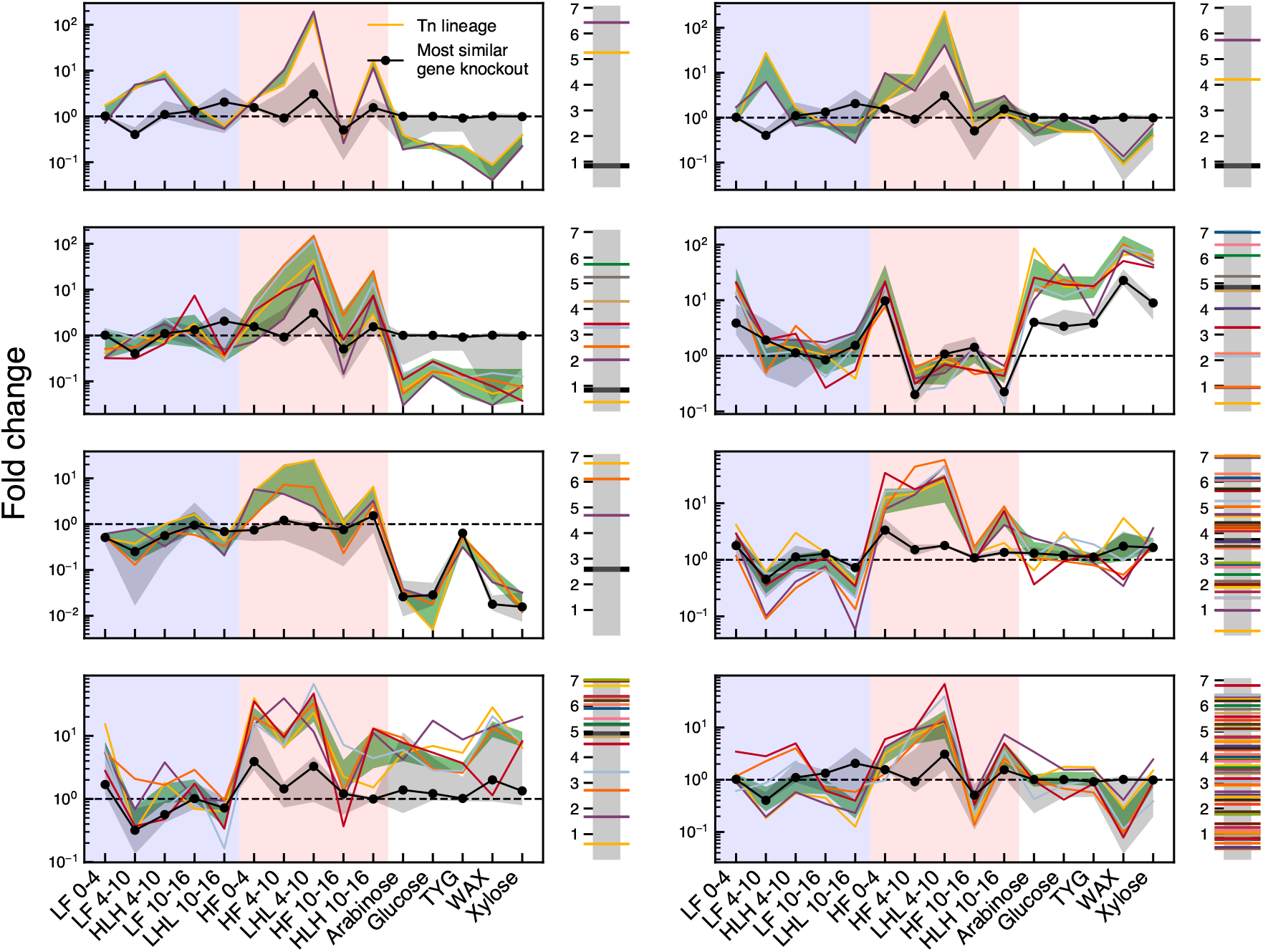
Analogous version of Fig. 4C,D. *Bc* lineages with similar fitness profiles(colored curves and green band; SI Section 6) that are dissimilar to any gene knockout (black) in the library.

**Figure S15:**
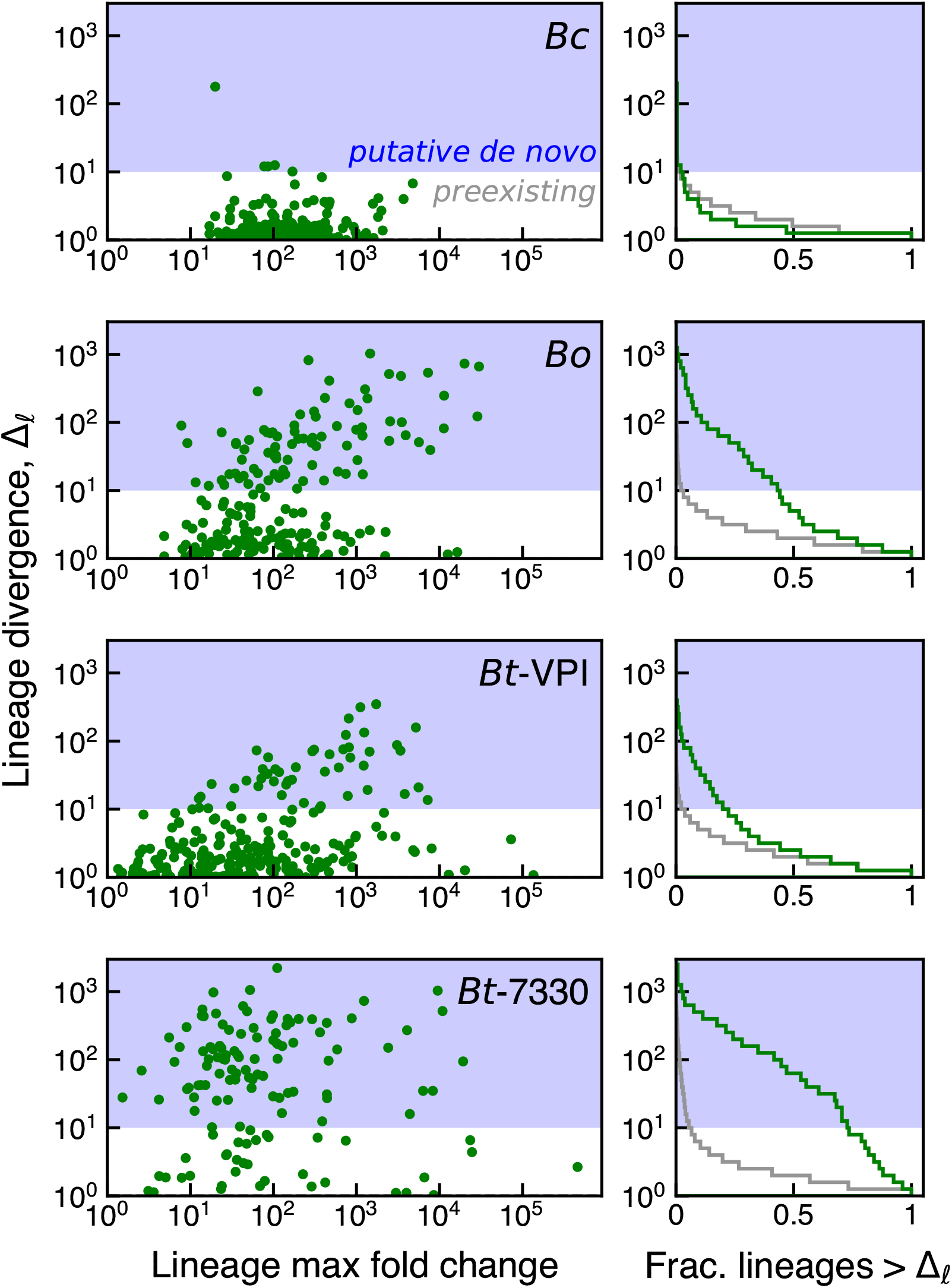
Evidence for adaptive *de novo* mutations from the variability across mice. **Left**: lineage divergence as defined by Eq. (32) (a measure of the variability of a lineage across 5 HF/HS mice, SI Section 7) as a function of the maximum fold change of that lineage in all mice from the same diet. Dozens of lineages that reached > 0.1% by day 16 in at least one mouse exhibited large divergences (> 10, blue shaded region) at this time point, suggestive of putative *de novo* mutations. **Right**: survival function (green curve) of the divergence metric for the lineages in the left plot. For comparison, the survival function of divergences for all lineages is shown in grey.

**Figure S16:**
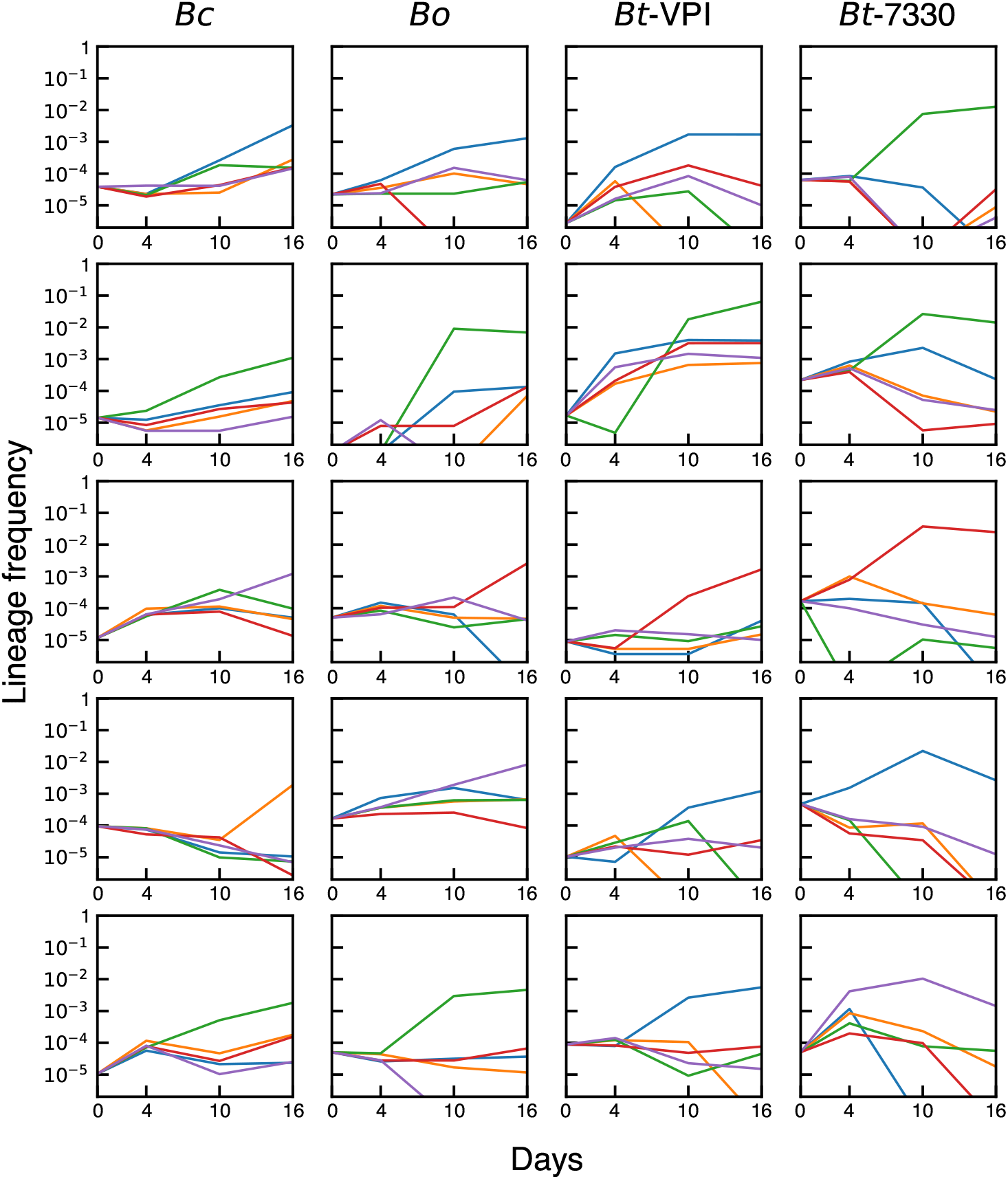
Trajectories of example lineages with putative *de novo* mutations. Five lineages with divergence >10 in each species (column) are shown. Lineages were randomly sampled from each of the three species (*Bo*, *Bt-VPI*, *Bt-7330*) in Fig. S15 that exhibited large numbers of strongly diverging lineages. Each lineage is illustrated in a separate panel. The individual curves represent the trajectories of that lineage in each of the 5 HF/HS mice. Each mouse is represented by the same color across panels.

**Figure S17:**
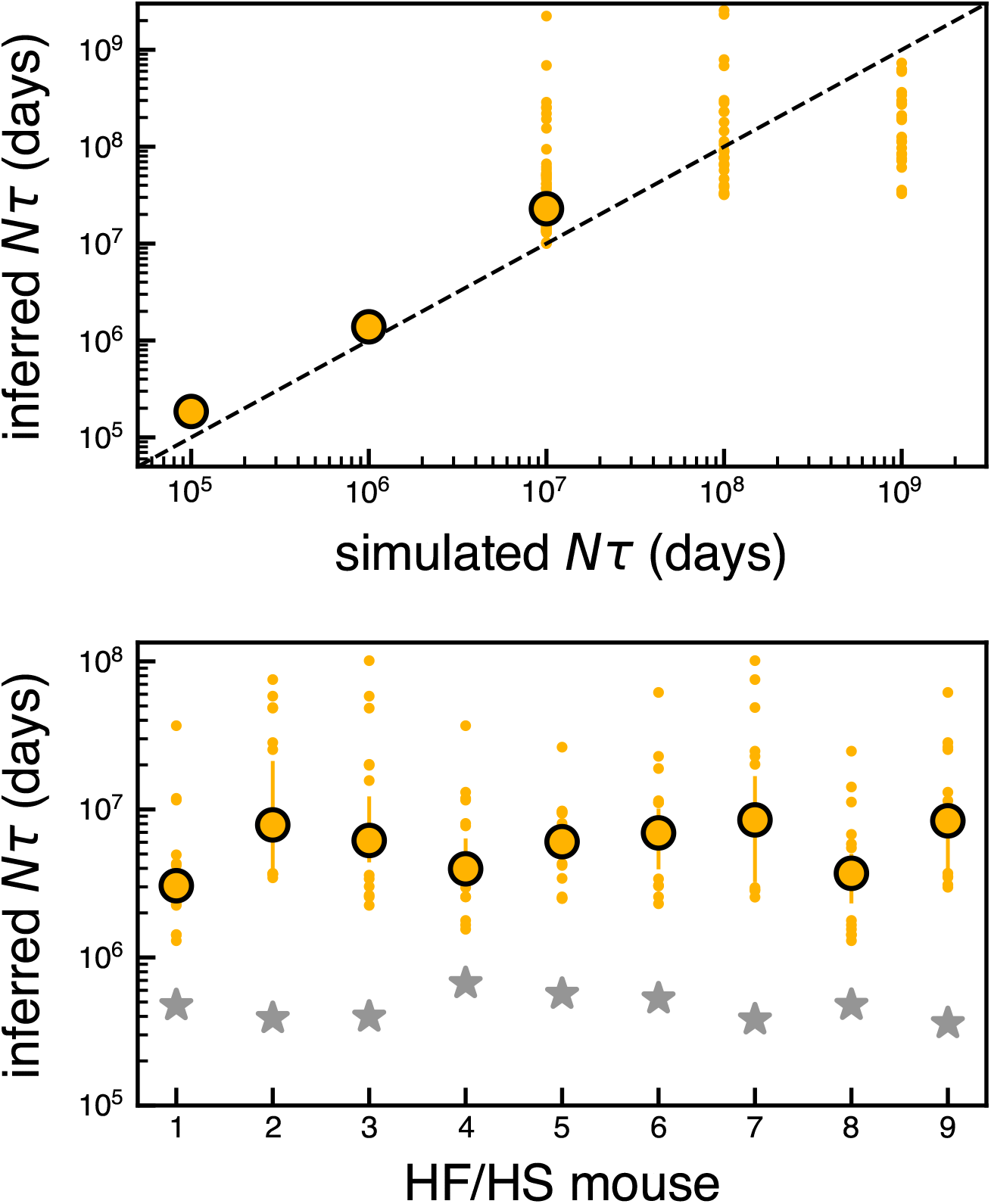
Estimating the strength of genetic drift in *Bc* during days 0-4. Inferred values of *N_e_τ_e_* from simulated **(top)** and empirical **(bottom)** *Bc* populations in HF/HS mice (SI Section 8). For both simulated and experimental data, different inferences were generated by shuffling mice among the separate cohorts used to estimate *ȳ_L_*, 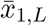, and 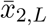 (Eq. (39)). The median (outlined circles), interquartile range (lines), and outliers (dots) of inferences are shown; when drift is weak (simulated *N_τ_* = 10^8^, 10^9^), the algorithm typically does not fit a finite *N_τ_*, so the median is not plotted. In the experimental data, a particular inference was assigned to a mouse if this mouse was used to estimate *ȳ_L_*. For comparison, the grey stars indicate sequencing depth at day 4.

